# Mathematical model of tumor-macrophage dynamics in glioma to advance myeloid-targeted therapies

**DOI:** 10.1101/2025.07.14.664717

**Authors:** Jesús J. Bosque, Jordan Martínez, José García Otero, Guim Aguadé-Gorgorió, Javier E. Sanchez-Galan, Juan Belmonte-Beitia

## Abstract

Recent biological research has highlighted the relevance of myeloid-cell populations in glioma growth, with a particular role played by tumor-associated macrophages (TAMs), which comprise resident microglia and monocyte-derived macrophages. Additionally, radiation therapy, the most common treatment for gliomas, significantly alters the tumor microenvironment, affecting TAMs and contributing to tumor recurrence. Promising preclinical studies have identified and developed drugs targeting TAMs. The development and combined deployment of these therapies require in silico techniques that enable us to optimize their outcomes. To do so, we need mathematical models of glioma growth and therapy response that explicitly incorporate TAMs—an often overlooked component in existing models. Here, we present a dynamical model of glioma growth driven by tumor-immune interactions. The model was parametrized using published data from mice experiments, including responses to ionizing radiation. We used this model to investigate glioma progression under radiotherapy combined with three treatments targeting distinct aspects of TAM biology. Simulations revealed that anti-CD47 enhanced the otherwise weak phagocytic activity, extending the upper tail of the survival curve. *α*-CD49d, which limits monocyte trafficking after irradiation, offered consistent survival benefits across digital twins of mice. Finally, CSF-1R inhibitors, which block the primary growth factor regulating TAM function, resulted in the largest overall survival improvement in silico. Our results aligned well with experimental evidence, suggesting that the model may help inform the optimization of myeloid cell-targeted immunotherapies, including their timing, dosage, and combination with radiation therapy, with potential relevance for improving glioma treatment strategies.

## 1. Introduction

Myeloid cell-targeted immunotherapies have the potential to revolutionize cancer treatment by addressing the most abundant immune population in the tumor microenvironment (TME) across many cancers [1]. While the past two decades have been dominated by the rise of T cell-targeted therapies, such as PD-1 and PD-L1 checkpoint inhibitors, which have achieved significant success in multiple cancers, myeloid cells remain a largely untapped target [2]. These cells not only represent the predominant immune population in most cancers but also act as key orchestrators of tumor progression and immunosuppression [3]. Emerging preclinical and early clinical studies demonstrate that myeloid cells are actionable targets and suggest interventions that can reprogram the TME to inhibit cancer progression and enhance immune-mediated tumor destruction [4]. As a result, myeloid cell-targeted therapies are poised to drive the next wave of advancements in cancer immunotherapy [5]. Gliomas are primary tumors arising in the central nervous system—normally, the brain—due to the abnormal proliferation of glial cells. Their treatment remains highly challenging, with poor survival rates, particularly in the case of glioblastoma (GBM). This has driven significant interest in developing new therapeutic approaches to improve patient outcomes [6]. Unlike other brain malignancies, such as brain metastases, which often exhibit high lymphocyte infiltration, gliomas are immunologically cold tumors. Instead, they are heavily infiltrated by myeloid cells, particularly macrophages, which can constitute up to 50 % of the total tumor cell population [7]. This distinct immune composition makes gliomas ideal candidates for therapies targeting myeloid cells, and specifically macrophages, to reshape the TME and enhance treatment efficacy [8].

Tumor-associated macrophages (TAMs) within the glioma TME originate from two distinct sources [9]. The first source are microglia (MG), which are brain-resident macrophages derived during embryonic development. The second source are monocyte-derived macrophages (MDMs), which infiltrate the tumor as circulating monocytes from the bloodstream and mature into macrophages after recruitment. Macrophages have traditionally been classified along a pro-inflammatory to anti-inflammatory axis, labeled as M1 and M2, respectively. While this binary classification has evolved to acknowledge the complexity and plasticity of macrophage biology, the terms “M1-like” and “M2-like” remain widely used in the literature [10]. In this context, M1-like TAMs exhibit anti-tumor activity, whereas M2-like TAMs promote tumor progression by supporting immunosuppression, angiogenesis, and tumor growth [11]. This dual behavior underscores the importance of understanding TAM phenotypes in glioma therapy by modeling their ecological interactions with tumor and other immune cells. Furthermore, tumor cells have the ability to hijack the global macrophage population to their own benefit, creating an anti-inflammatory, tumor-promoting habitat populated by M2-like TAMs [2, 12].

Ionizing radiation (IR) is a cornerstone of glioma treatment [13], effectively reducing tumor size, slowing progression, and alleviating symptoms. However, despite its relative effectiveness, IR is not curative in gliomas. The response to and resistance against IR are heavily influenced by the state of the TME [14]. On the other hand, IR significantly impacts the TME, such as by stimulating the infiltration MDMs, which can contribute to tumor regrowth and immunosuppression [15–18]. As TAM-targeted therapies are unlikely to be implemented as standalone treatments, it is essential to understand their interactions with IR. A combined approach leveraging TAM modulation alongside IR could optimize therapeutic outcomes by enhancing anti tumor activity while avoiding a tumor-promoting microenvironment. This synergy has the potential to address the limitations of each modality, offering more effective treatment options for glioma patients [19].

Mathematical modeling of tumor growth and treatment response is an invaluable tool for analyzing treatment strategies, optimizing therapies, and advancing personalized medicine to inform clinical decision making [20, 21]. The modeling of macrophages in the TME has received comparatively limited attention compared to its biological relevance. Early work focused on the anti-tumor action of macrophage-lymphocyte collaboration [22]. Pioneer studies by Owen and others included the co-evolution of tumor cells and TAMs [23–25]. Interest reignited slowly over the last decade [26, 27], eventually giving rise to a new wave of mathematical works on TAMs [28–34]. See ref. [35] for an exhaustive review.

Gliomas represent one of the most commonly studied tumors in the mathematical oncology literature [36–41]. However only a minimum number of modeling studies have considered the key role of TAMs in glioma development [42–44]. In [42], the authors developed a spatial model to study the interaction between glioma cells and TAMs through the EGF–CSF-1 signaling pathway, aiming to understand how this paracrine loop contributes to glioma progression. Simulation results highlighted the correlation of CSF-1 concentration with tumor grade. Similarly, ref. [43] used a reaction-diffusion model to study the interaction of glioblastoma cells with MG via signaling molecules like CSF-1, EGF, and TGF-*β*, showing that tumor-secreted CSF-1 activates MG to release TGF-*β*, which in turn enhances tumor invasion. Finally, ref. [44] demonstrated that TAM density at a glioblastoma biopsy location is a critical factor in predicting post-surgical recurrence, challenging standard biopsy approaches and highlighting TAMs as key drivers of recurrence. Beyond the intrinsic value of these articles, to our knowledge no modeling work has specifically accounted for interactions between TAMs and radiation therapy using real data and the effect of cutting-edge myeloid immunotherapies.

Here, we present a mathematical model of glioma growth that captures the ecological interactions between different cell types using ordinary differential equations (ODEs). The model incorporates the role of TAMs, accounting for their dual behavior in both sustaining and opposing tumor growth. We further extend the model to include the effects of IR on both tumor cells and immune populations and parametrize the tumor using published data from experimental gliomas in mice [45]. Finally, we use the model to simulate treatment scenarios involving myeloid-targeted immunotherapies, exploring their potential to modulate TAM behavior and enhance therapeutic outcomes. We believe this model can serve as a foundation for the development and optimization of immunotherapies, ultimately contributing to improved treatment strategies for gliomas.

## 2. Materials and methods

### 2.1. Mathematical model of glioma growth

To model the temporal dynamics of the main populations composing a glioma we use a system of three coupled ODEs. Each of these equations correspond to the number of **tumor cells** (*T*), **anti-tumor macrophages** (*A*), and **pro-tumor macrophages** (*P*), respectively. All three populations proliferate at rates *ρ*_*T*_, ρ_*P*_, and *ρ*_*A*_following a Gompertzian growth limited by maximum attainable tumor volume (carrying capacity *K*). The macrophage and tumor populations are embedded in a strong cooperative interaction, in which both cell types proliferate in the presence of the other [12]. The macrophage population proliferates only in the presence of tumor cells, while the tumor cells, in turn, divide only when they receive the appropriate signals from pro-tumor macrophages.

Initially, the microglia naturally present in the brain have an anti-tumor behavior, killing tumor cells at a rate *µ*. The macrophage population *A* is therefore composed exclusively of microglia. However, tumors have the ability to hijack microglia activity to turn it to a pro-tumor phenotype [12]. Therefore, there is a transfer of anti-tumor microglia (*A*) to the pro-tumor population (*P*), which happens in the presence of tumor cells with a rate *λ*. There is only a certain number of microglia that can turn into the pro-tumor phenotype, therefore, this process is limited by a Michaelis-Menten term with the constant *K*_*λ*_. The invasion of new areas in the healthy brain by diffusing tumor cells recruits more microglia, which subsequently become part of the tumor (term 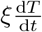).

Pro-tumor macrophages, in turn, have two different sources. One is the direct transformation of microglia mentioned in the previous paragraph, while the second is the infiltration of MDMs from the vasculature triggered by the presence of tumor cells [9]. The latter is modeled here by a flux term *u* (fig. 1) that is directly proportional to the existing number of tumor cells (*ϕ·T*). All the infiltrating MDM gathered by the tumor cells are assumed here to exhibit a pro-tumor behavior. The population of pro-tumor macrophages described is therefore composed of a mixture of MG and MDM. Finally, both pro-tumor and anti-tumor macrophages have a limited lifespan, which has been modeled by a decay with a time constant *τ*_*M*_.

**Figure 1.**
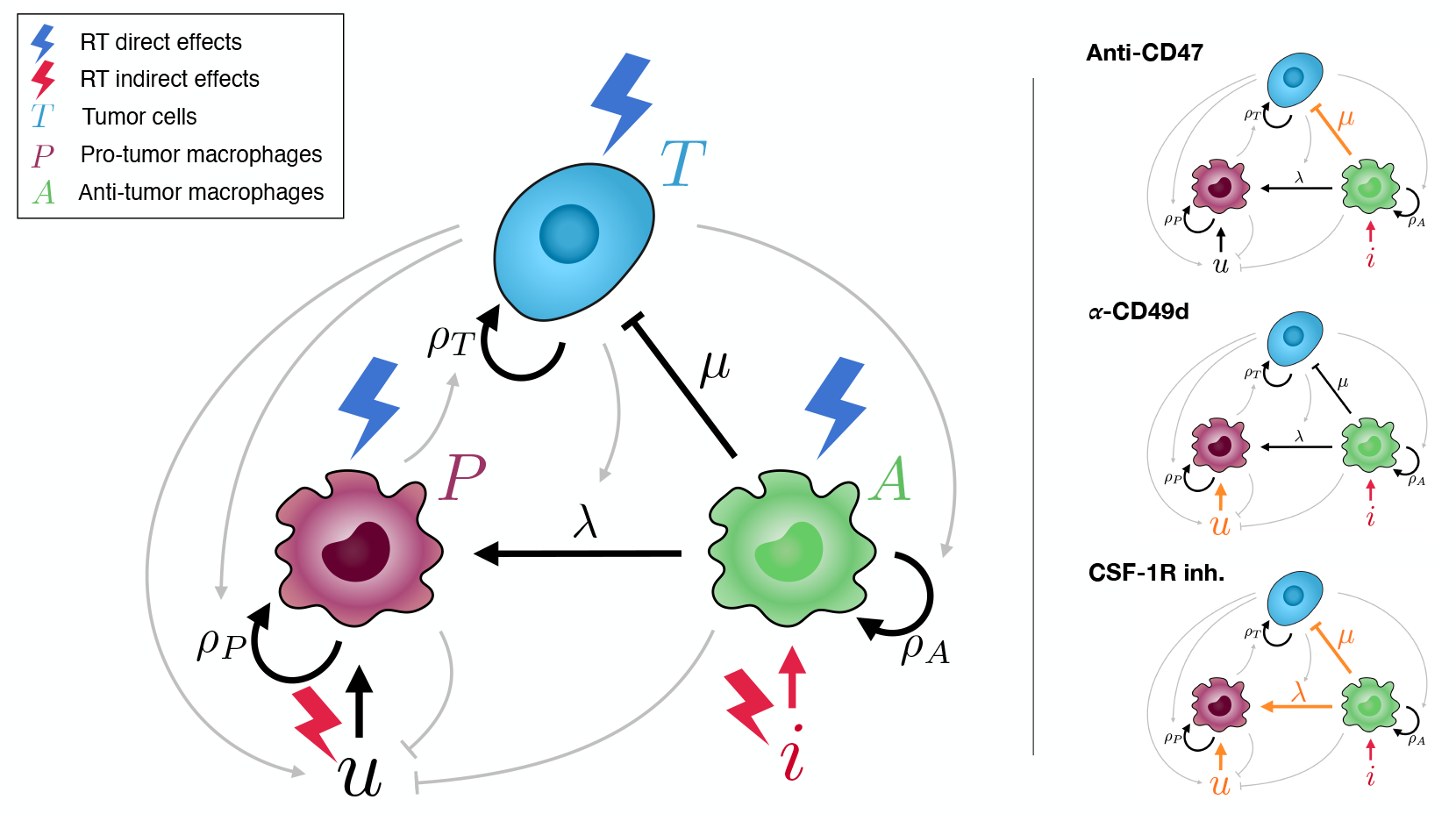
Diagram of the model describing the network of interactions between cells. Initially, resident microglia constitute the antitumor population that kill tumor cells at a rate *µ*. The presence of tumor cells promotes their change to a pro-tumor phenotype, which in turn promote tumor growth, at a rate proportional to *λ* and the number of tumor cells. Tumor cells also recruit myeloid derived macrophages (MDMs) from the vasculature (*u*) which have a pro-tumor behavior. The application of radiation therapy reduces the amount of every population, but also triggers inflammation (*i*) and increases the flux of MDMs through induced damaged in the vasculature. The left hand side panel illustrates the links affected by the three types of myeloid cell-targeted immunotherapies considered here.

The previous considerations lead to the following system of differential equations

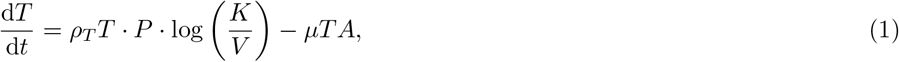

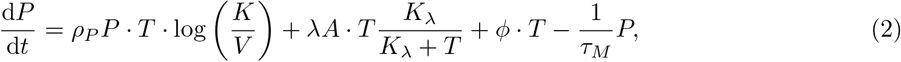

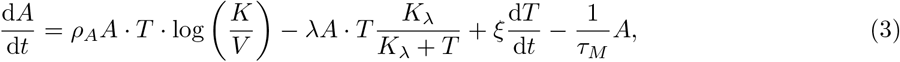

where the total volume of the tumor is the sum of the different populations weighted by their respective unitary volumes, *v*_*T*_and *v*_*M*_

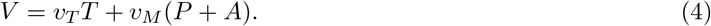

Additional considerations regarding the assumptions made here and their justification are included in the Supplementary Information (SI), in SI1 “Assumptions of the mathematical model”. For the numerical resolution of the system we additionally tracked the contributions to *P* of MG and MDM individually (see SI2 “Individual tracking of MG and MDM”).

### 2.2. Modeling the effect of ionizing radiation

The effect of radiation on the populations is primarily modeled as an instantaneous damage that transfers a fraction of the populations (1*−S*) to populations of damaged tumor cells (*T*_*D*_) and damaged macrophages (*M*_*D*_)

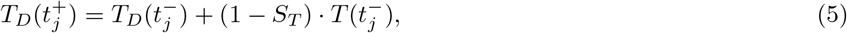

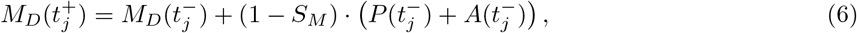

where *S*_*T*_and *S*_*M*_are the survival fractions (between 0 and 1) of tumor cells and macrophages after therapy, and *t*_*j*_ are the times at which different fractions *j* of IR are administered. This damage is indicated in fig. 1 by blue lightning symbols pointing at the populations.

The damaged populations contribute to tumor volume after therapy,

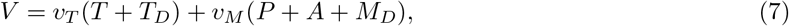

but they do not exhibit any of the functional behaviors of their undamaged counterparts. They are part of the tumor until damage becomes irreparable and cells die with characteristic times of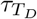and 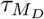. Thus, their dynamics are described by

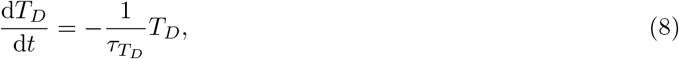

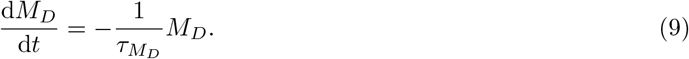

Moreover, the effect of radiation causes secondary effects that modify the TME (crimson lightning symbols in fig. 1). First, IR causes an increase in MDM infiltration [15, 16], increasing the flux *u* of external MDM. Similar to eqs. (5) and (6), this increase in the flux is modeled as an increase in the parameter governing the dependence of MDM influx on the number of tumor cells

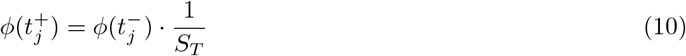

Notice that the parameter *ϕ* is constant along the simulation, with the exception of changes occurring instantly after exposure to IR. In this work we assumed that the increase in MDM influx is proportional to the therapeutic efficacy of IR (the damage affecting tumor cells), using the inverse of the survival fraction *S*_*T*_as a measure of MDM flux increase.

In addition, IR induces inflammation in the TME, harnessing local microglia to help clear dead cells and debris [12]. We modeled this inflammation by adding an additional flux *i* to the rate of change of anti-tumor macrophages proportional to the total number of damaged cells (*T*_*D*_+ *M*_*D*_). Therefore, after exposure to IR, the equation that governs the progression of anti-tumor macrophages is

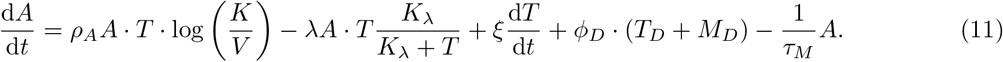

The way of integrating these effects in the simulation algorithm is that the numerical resolution stops at the time of radiation treatment and then restarts with new initial conditions based on the current cell counts and the survival fraction. With the mechanisms described in this subsection, we cover the effect of radiation therapy both as a direct killing of cells and effects on the immune microenvironment.

### 2.3. Experimental data

We contrasted and parameterized our model with empirical data from previously published experiments using PDG-Ink4a/ArfKO immunocompetent mice [45]. We used volumetric follow-up data from 4 untreated mice and 10 mice treated with IR. Additionally, we used bulk data from the markers Ki67^+^ (cell proliferation), Iba1^+^ (macrophages), and CC3^+^ (apoptotic cells) at four time points: control, after 5 days of IR (5 d), 21 days after the initiation of IR (21 d), and at tumor recurrence (IR Rec). Finally, the percentages of MG and MDM had been measured by CD49d^−^(MG) and CD49d^+^ (MDMs) flow cytometry markers. See SI3 ‘Data from experiments in mice’ for a detailed explanation of the experimental data used. These biomarker data were compared to the percentage variables macrophages, MG, MDM, proliferation, apoptosis, and killing, all calculated from the mathematical model as described in SI4 “Calculation of markers from the model”. Throughout the figures, unfilled markers represent metrics from in silico simulations, and filled markers represent experimental data.

### 2.4. Numerical resolution

#### 2.4.1. Control cases

To simulate the growth of untreated tumors, we solved the system eqs. (1) to (3) numerically using the routine ode45 of MATLAB, version R2023b. Each of the temporal dynamics for the four control experimental mice available in ref. [45] was reproduced in silico. To do this, we used the volume at the first experimental point as initial condition for *V*, eq. (4). This volume is the sum of individual volumes of each population and therefore must be divided among them to assign initial conditions for each equation. The fractions corresponding to each population are approximately known from the original data reference: in the untreated control the fraction of tumor cells is approximately *f*_*T*_= 0.96, the fraction of MG (assumed to be entirely anti-tumor initially) is *f*_*A*_= 0.035 and the fraction of MDM, *f*_*P*_= 0.005. The initial volume can then be distributed among each of the populations as

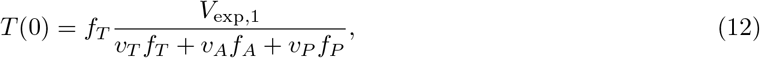

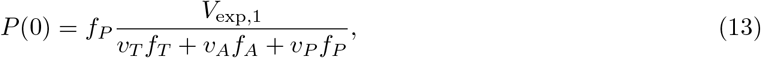

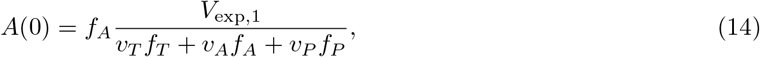

where *V*_exp,1_ is the volume at the first experimental time point. For the individual tracking of MDM and MG, the initial number of pro-tumor macrophages at the initial time is divided equally between the two subpopulations *P*_1_ and *P*_2_.

The simulations for each individual used the same estimated parameters, except for the proliferation rate of the tumor cells *ρ*_*T*_, which was used as a single free parameter to fit to the volume data at the available data points.

#### 2.4.2. Treated cases

The simulations corresponding to mice treated with IR were solved numerically in a piecewise manner using MATLAB’s `ode45` during continuous intervals without treatment. IR was modeled as described in section 2.2 by changing the proportions of the respective populations at the treatment times and using those new initial conditions for the next interval.

The initial conditions prior to the IR applications were set similarly to those for the control cases described in the previous section 2.4.1. However, there are differences in the volume values between both datasets. While the control mice had initial tumor volumes around 3 mm^3^, the initial volumes for the treated cases were much larger (mean: 41.5 mm^3^). This suggests that the proportions experimentally available for control cases must be adapted to more advanced stages of growth.

To do this, we first determined the dynamics of the proportions of different populations in the previously fitted control cases. Next, we related these proportions to the tumor volume at each time point, expressed as a fraction of the maximum volume reached. Using linear regression, we established relationships between the relative tumor volume at a given time and the proportions of each population. For a treated case with a specific initial volume, we calculated the ratio of that volume to the maximum volume observed in the corresponding control case. This ratio was then used as input for the linear regression models. The outputs of the regressions provided the population proportions, which we used to divide the tumor volume among the respective populations in the treated cases.

The simulations for each individual case are then run following the model explained previously, using the first experimental volume to set the initial condition in the simulations. In the IR cases, two free parameters are used to fit the individual cases: the proliferation of tumor cells *ρ*_*T*_, as in the control cases, and additionally the survival fraction of tumor cells *S*_*T*_, used as a metric of treatment effectiveness.

The remaining parameters used in the simulations are the same as in the control cases, with the exception of the tumor cell killing rate by anti-tumor macrophages *µ*. The implications of this change are discussed below.

### 2.5. Parametrization

To reproduce the temporal dynamics of tumor volume in the control cases we fixed all the parameters except the cell proliferation rate *ρ*_*T*_, which is considered to be individual for each subject and is used to fit the data. To perform this fit, we used the `fminsearch`routine from MATLAB to find the parameter minimizing the root mean squared error (RMSE) between the volume resulting from the model and the *n* volume data points *V*_*exp,k*_

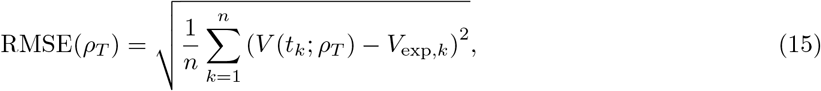

where *V* is the volume returned by the model as described by eq. (4).

In the same way, each case in the IR dataset was also fitted by minimizing the RMSE. In this case, two parameters were used: one ruling the overall tumor growth behavior, i.e., the tumor proliferation rate *ρ*_*T*_, and another accounting for the efficacy of the treatment, the tumor cell survival fraction *S*_*T*_

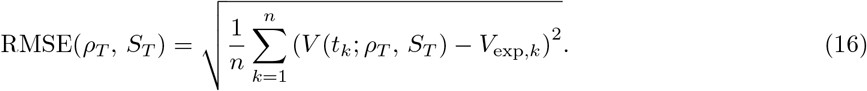

The availability of individualized experimental subjects with time follow-up allowed us to fit each case using the previous method.

With the exception of some easily inferable parameters (e.g. *v*_*t*_, *K*), the values of most fixed parameters are difficult to estimate. Due to the non-linear interactions between cell types existing in our model, literature values are not available and a full parametrization would require dedicated experiments.

To estimate these parameters we compared the assembled information provided by the experimental biomarker measurements to the analogous metrics given by our model (eqs. (S4) and eqs. (S6-S9) from SI4 “Calculation of markers from the model”). Because the experimental metrics are not linked to individualized subjects, both experimental and simulated data were treated at the group level.

We performed repeated fits of the individual cases’ time evolution as explained above for different sets of the fixed parameters. For each set we assessed volumetric fits and compared the trends in simulated biomarkers to those observed in the experimental setting. We iteratively modified the values of the fixed parameters manually to improve qualitative agreement with experimental biomarkers data, which is feasible thanks to the biological interpretability of these parameters.

### 2.6. Statistical analyses

To compare biomarkers values between groups of individuals undergoing different treatment modalities we calculated effect size using Cohen’s d along with each confidence interval (CI). We used MATLAB’s meanEffectSize for this purpose. When the confidence interval contained the value zero we labeled the effect as “inconclusive” using the mark abbreviation inc. The computation of Cohen’s d considers the difference between the mean of the first group and the mean of the second, therefore, a positive sign implies that the first group has a bigger mean, while a negative sign means that the second group’s mean is larger.

On top of the data values for the individual cases, each figure depicts an error bar showing means *±* standard error of the mean (SEM); this follows the standard in biology intending a direct translation from our figures to the main reference from which we gathered experimental data.

To compare survival differences between groups subjected to distinct interventions, we used Kaplan Meier curves. Statistical significance in survival differences were assessed using the log-rank tests, which compare the observed and expected number of events in each group over time under the null hypothesis of equal survival functions. Additionally, graphs show lower and upper confidence confidence bounds for the evaluated function values at a 95 % confidence level using dashed lines.

A linear regression was used to identify correlation between survival benefits in each of the therapeutic interventions and the fitted proliferation rate *ρ*_*T*_of each mouse. We also used receiver operating characteristic (ROC) curves to analyze the power of the proliferation rate as a binary classifier in the high and low benefit groups. We reported the area under the curve (AUC) in the ROC curve as a measure of how well the proliferation parameter separates the two groups. We used MATLAB’s function `fitlm` to fit the linear models, and `perfcurve` to obtain the ROC curve statistics for the binary classification.

## 3. Results

This section is structured as follows: (1) Adequacy of the model to reproduce experimental data (sections 3.1 and 3.2). (2) Mice digital twins upon radiotherapy treatment (sections 3.3 to 3.5). (3) Macrophage activity and anti-CD47 treatment (sections 3.6 and 3.7). (4) Treatment with *α*-CD49d (section 3.8). (5) CSF 1R inhibition (section 3.9).

### 3.1. The model is consistent with the volumetric data from experiments

We first proved the existence, uniqueness, and positivity of the solution of the system, and identified the system’s equilibrium points along with their stability (see SI5 “Basic properties of the mathematical model”). Subsequently, we analyzed the sensitivity to changes in the parameter values (SI6 “Sensitivity Analysis”).

Having demonstrated the mathematical suitability of our model to describe the dynamics of the studied populations, we sought to determine whether its solutions could accurately represent the real dynamics of experimental gliomas. To this end, we utilized previously published data on the growth dynamics of untreated gliomas in PDG-Ink4a/ArfKO mice (see section 2.3 and SI3 “Data from experiments in mice”), which included the volume data of four specimens. We computed the volume predicted by the ODE model using eq. (4) and fitted the data with a single free parameter for each animal, as described in section 2.5. We chose the proliferation rate of tumor cells, *ρ*_*T*_, as the free parameter for several reasons. First, its significance was highlighted in the sensitivity analysis, which identified it as the primary parameter influencing temporal dynamics (SI6 “Sensitivity Analysis”). Second, given the biological variability that affects all parameters, we hypothesized that the greatest variability among experimental individuals arises from heterogeneity in tumor cell behavior. In contrast, other system components, particularly those influencing macrophage behavior, are expected to exhibit greater homogeneity.

The model was able to fit the progression data observed in the untreated data well using *ρ*_*T*_as a single free parameter (fig. 2A). Nevertheless, a larger number of time points would be desirable to better assess the quality of the fit. Next, we investigated whether the dynamical model, combined with a simple model of IR effects (see section 2.2), could also reproduce the temporal dynamics of tumor volumes in PDG Ink4a/ArfKO mice undergoing radiation treatment. In addition to the parameter representing inter-subject heterogeneity (*ρ*_*T*_), we introduced a second free parameter to account for individual differences in response to IR, specifically radiation sensitivity. For this, we used the survival fraction of tumor cells, *S*_*T*_, as the second free parameter. Data from 12 mice were available; two were excluded because they only included two time points. The remaining ten, with 4 to 7 time points each, were fitted using *ρ*_*T*_and *S*_*T*_as free parameters. For 8 out of the 10 specimens, we observed good agreement between the volumetric data and the model fits, (fig. 2(B)). Two cases did not show a good fit to the data, even though the general trends were captured. Interestingly, both cases exhibited a notable tumor decrease during two consecutive time points. These cases warrant further investigation to determine whether the model encounters challenges in representing tumor responses over prolonged periods after treatment.

**Figure 2.**
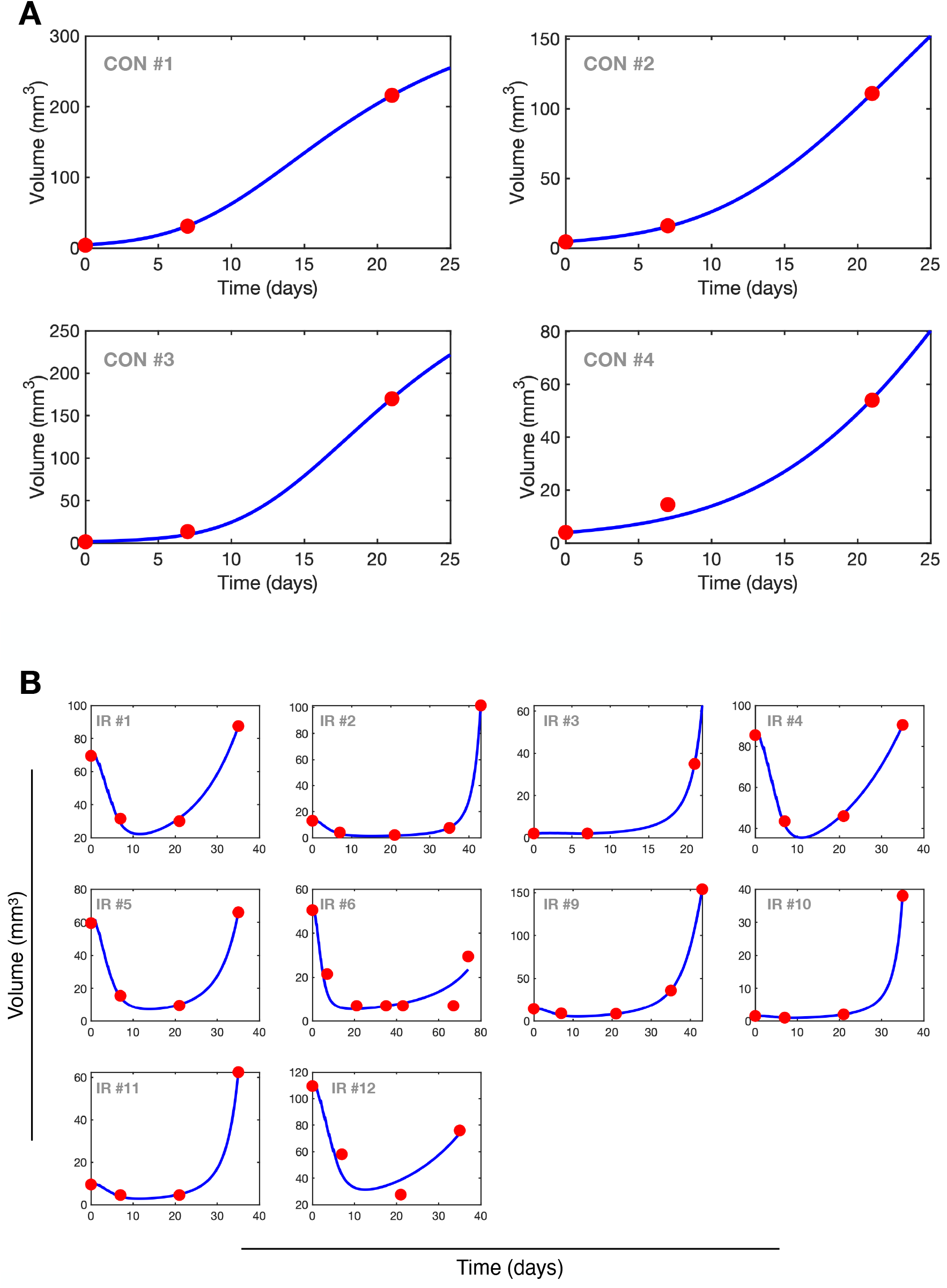
Tumor volume dynamics. Red dots represent the available data on tumor growth in PDG-Ink4a/ArfKO mice for control cases (A) and mice treated with five daily fractions of IR between days 1 and 5 (B). The blue line corresponds to the tumor volume predicted by our model, fitted to the data using the proliferation rate of tumor cells, *ρ*_*T*_, as a free parameter for control cases, and both *ρ*_*T*_and the IR survival fraction, *S*_*T*_, as free parameters for treated cases.

Notably, all fixed parameters used to fit the control and treated cases were the same, except for the tumor killing rate exerted by antitumor macrophages, *µ*. To match the observed dynamics, we found that significantly different values were required between the treated and untreated cases (*µ* = 1.2 *×* 10^−4^d^−1^ cell^−1^for the control group and *µ* = 5 *×* 10^−7^d^−1^cell^−1^for the IR group). We hypothesize that this discrepancy may reflect actual biological differences between the two groups. Specifically, the untreated cases exhibited very small tumor volumes at early time points compared to the treated cases, which had grown much larger over a similar period. Consequently, the control group may demonstrate a greater ability of the antitumor macrophages in the TME to oppose tumor growth.

Altogether, these results demonstrate that our model can describe the temporal dynamics of both untreated experimental gliomas and gliomas treated with IR using a minimal set of parameters (1 and 2, respectively).

### 3.2. The model captures the trends in biomarker changes

In addition to the experimental data on the longitudinal growth of tumor volumes, ref. [45] included measurements for three key biomarkers: the percentage of macrophages, the amount of apoptotic cells, and the proportion of proliferating cells (see SI4 “Calculation of markers from the model”). These experimental measurements are shown on the right-hand side of each panel in fig. 3, using filled markers. To further validate our mathematical model, we computed the equivalent variables from our simulations using eqs. (S4), (S8), and (S9) from the SI to calculate the corresponding biomarker in silico values (unfilled markers in the left-hand side panels of fig. 3). The model qualitatively reproduced the changes observed in these biomarkers, including: (i) a sharp increase in macrophage population immediately after IR treatment, followed by a return to baseline levels at long times and recurrence (fig. 3(A)); (ii) an increase in the number of apoptotic cells after treatment, with the model predicting a return to baseline far from the treatment period (fig. 3(B)—no experimental data for 21 d and IR Rec); and (iii) a decrease in proliferation after IR, with much higher proliferation rates at tumor maturity and recurrence (fig. 3(C)).

**Figure 3.**
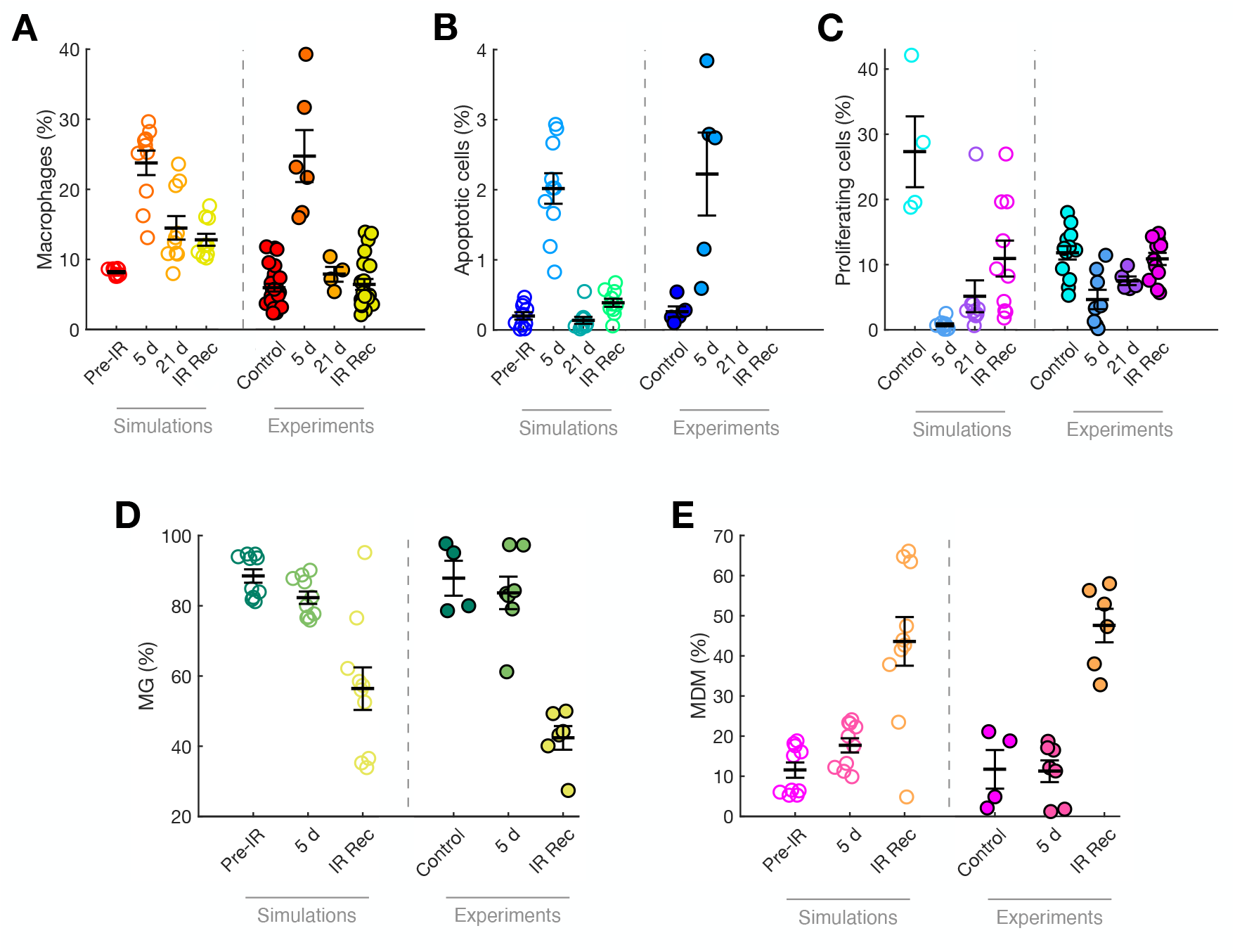
Changes in tumor biomarkers in simulations and experiments. The left hand side of each panel shows the measurement resulting from our model using eqs. (S4), (S8) and (S9) (open markers). The right hand side shows the values obtained experimentally in PDG-Ink4a/ArfKO mice (ref. [45]) for (A) Iba1^+^ cells, (B) CC3^+^ cells, (C) Ki67^+^ glioma cells, and flow cytometry quantitation of (D) CD49d^−^MG and (E) CD49d^+^ MDMs (closed markers). Graphs show means *±* SEM.

Additionally, the distribution of macrophages between MG and MDM had been measured experimentally, providing a good overview of how these populations change after IR. Our model showed excellent agreement with the experiments, indicating that shortly after IR, the population is still dominated by MG, while recurrence is characterized by a significant decrease in MG numbers and a shift towards dominance by infiltrating MDM.

It should be noted that traditional fitting is not possible for these data, as individual cases were not identified and there is no individual follow-up. Therefore, while the qualitative agreement is not perfect, it supports the validity of the model. We have deliberately omitted measures of effect size and statistical significance in fig. 3, as our aim is not to examine specific pairwise group differences or assess the extent to which experimental findings are replicated across groups, but rather to illustrate the qualitative similarity between the simulated and experimental systems.

Altogether, both the data and the model show that IR induces an increase of the number of macrophages inmediately after the treatment and that recurrence is associated with both an increase in proliferation and a shift toward infiltrating MDM in the distribution of TAMs.

### 3.3. Temporal dynamics of the model variables

Our model enables the identification of the different populations driving overall tumor growth. We tracked the temporal dynamics of the three main populations—glioma cells (*T*), anti-tumor macrophages (*A*), and pro-tumor macrophages (*P*)—in the fitted control cases. The results reveal a steady exponential growth in the tumor population *T*, which reaches a limitation at high numbers in the first three cases (fig. S4(A)), but not the fourth due to its smaller size. The macrophage population in the untreated cases follows the growth of tumor cells, with a progressive dominance of pro-tumor macrophages (*P*) over their anti-tumor counterparts (*A*).

Regarding the simulation of treated cases (fig. S4(B)), simulations show the reductions mainly in the tumor population, due to the effect of IR, but also in the immune populations. Note that the nonlinear interactions described by the model lead to divergent behaviors, with pro-tumor macrophages either increasing or decreasing depending on the case. The effect of IR is followed by a regrowth in all populations that shows evidence to be accelerated, specially in the tumor population *T*. This acceleration post-treatment would be an effect of the positive feedback loop existing between *P* and *T*.

In terms of the distribution of MG and MDM, the control cases show an steady distribution that is only very lightly affected by time (fig. S5(A)). Even though there is a slight tendency to increase the number of MDM, the proportion of MG and MDM persist almost constant during untreated tumor growth. In contrast, this proportion is highly disrupted in the treated cases (fig. S5(B)), where IR is followed by a steady increase in the number of MDM at the expense of MG. In some cases, this is reverted at long times due to the fast pace of growth of tumor growth and the inclusion of new MG due to the term *ξ* d*T /* d*t*.

### 3.4. Radiation therapy survival benefit

Treatment with IR is one of the primary modalities used against gliomas. We sought to determine the survival benefits generated by IR treatments for the treated PDG-Ink4a/ArfKO mice using our model. To evaluate the effectiveness of IR within our in silico framework, we created the following two cohorts:

- **Digital twins from experiments:** The IR treated cohort consisted of parametrizations of the treated specimens obtained by fitting the real experimental data as shown in the previous subsection (fig. 2(B)).
- **Untreated version of the treated digital twins:** For each digital twin (i.e., the combination of the model and individual parametrizations) generated from the experimental cases treated with IR, we simulated an alternative scenario by removing any treatment. This allowed us to observe the tumor progression in each particular digital twin without IR intervention.

As a criterion for the death of the digital twins in both cohorts, we set a volume threshold consisting of the volume value at the last experimental point increased by 20 %. The time at which the simulated tumor exceeded this volume was taken as the time of death. We chose this criterion over previously employed ones, like a fixed constant fatal tumor volume [46], or a randomly assigned variation around a fixed threshold [47], in order to maximize the use of individualized information from the experiments with mice, namely that the tumor volume at death for each mouse was at least that of the last observation.

We ran each simulation for the treated and untreated digital twins and compared the survival times of both cohorts, as shown in the Kaplan-Meier curve (fig. 4). This quantifies the benefit of the model in assessing the effect of IR. The log-rank resulting p-value was 0.006, indicating a significant difference between the survival curves of the treated and untreated cohorts. The median survival gain attributed to IR treatment was 18 days. Notably, data on the survival benefit of radiation therapy in the studied mice are available (see Ref. [45] and the inset in fig. 4). The experimental treatment also resulted in a significant difference between cohorts, with a median survival benefit of 18 days. Therefore, our model accurately captures the effects of radiation therapy on PDG-Ink4a/ArfKO mice and the associated survival benefit.

**Figure 4.**
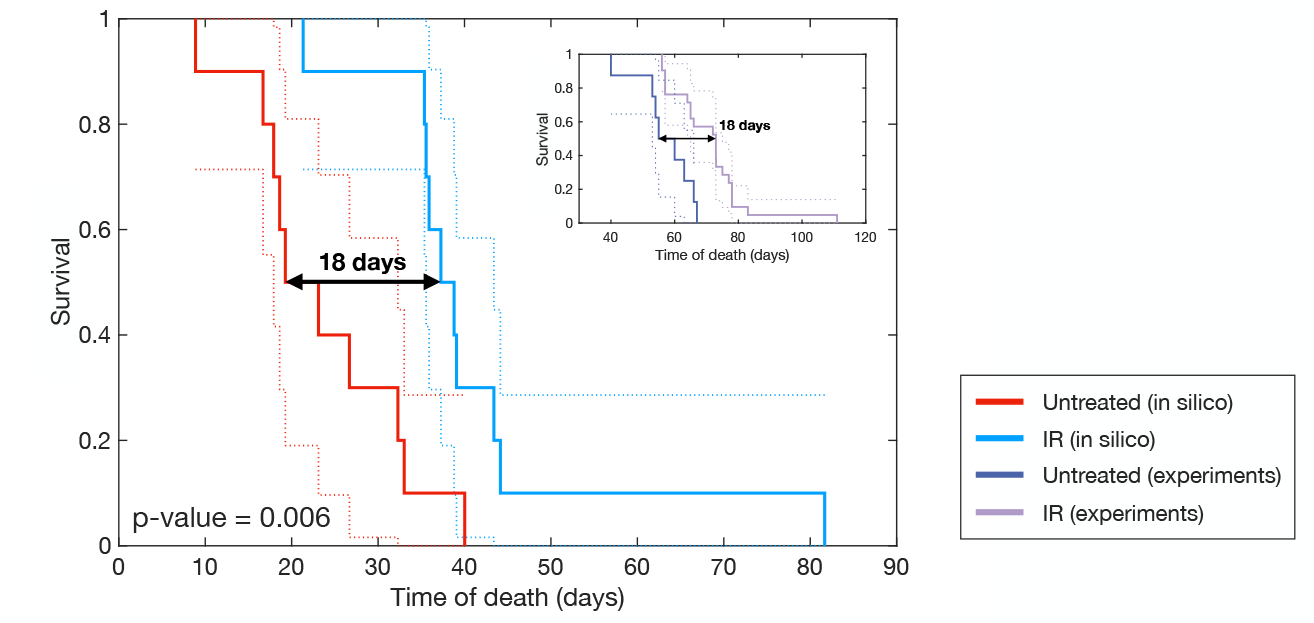
Survival benefit of IR in mice digital twins. Kaplan-Meier curves showing the survival difference between untreated mice digital twins (red) and mice treated with 5 daily fractions of radiation therapy (blue). IR results in a median survival benefit of 18 days, with significant survival difference (log-rank p-value = 0.006). The inset displays the equivalent experimental results taken from ref. [45].

### 3.5. Protracted radiation therapy provides a modest survival benefit

Previous research has suggested that spacing out the time between TMZ doses [47, 48] and radiation therapy fractions [49, 50] could enhance the survival benefit provided by both therapies. As discussed above, the effect of IR not only reduces tumor volume but also induces an increase in MDM, which are linked to glioma recurrence. This led us to question whether spacing out IR fractions (a treatment modality known as *protracted* radiotherapy) could delay MDM infiltration and improve the treatment response.

To evaluate this, we conducted simulations of the treated digital twins undergoing treatment with five IR fractions separated by different time intervals. For each fractionation, we analyzed the survival difference compared to the standard protocol at three group segmentations: median survival, the 75th percentile of survival (good survival), and the 25th percentile of survival (poor survival).

We found that median survival improves for fractionation intervals greater than 2 days, reaching an improvement up to nearly 9 days of interval, for which the benefit rapidly drops (fig. S6(A)). However, this gain is small (less than 3 days of survival benefit for the maximum protraction) and is unlikely to be detected in a randomized clinical trial. Moreover, spacing out the treatment reduces the benefit of tumor volume reduction: more spaced fractionations, while offering some survival benefit, result in larger tumor volumes, with this effect increasing as the fractionation spacing widens (fig. S6(A), inset). Consequently, gliomas in the 25th survival percentile do not benefit from protraction; instead, they fare worse than with the standard protocol. This aligns with previous research, where spaced TMZ protocols were effective for low-proliferating gliomas but not for more aggressive ones [48]. In the most favorable cases, the fractionation also provides a slight benefit, though this gain is smaller than for median survival gliomas and nearly imperceptible.

Another way to look at this results is representing the survival benefit as a color variable in a map formed by each individual mouse (x-axis) and the increasing interval between fractions (y-axis). This is represented in fig. S6(B), which shows that some in silico mice get an increasing survival benefit from IR protraction, while for other there is a decrease in survival.

We then wondered whether the fitted proliferation rate *ρ*_*T*_could classify the individuals benefiting from IR protraction. We found that for protracted IR with a 8 days gap between consecutive IR fractions there is a good fit (*R*^2^ = 0.76, F-test p-value *<* 0.001; see fig. S9(A)) between the survival benefit and the logarithm of the proliferation rate. This means that mice with lower proliferation rates are more likely to benefit from IR protraction. Actually, six out of the ten mice have a positive survival benefit, with four mice experiencing an earlier death under the application of protracted IR. We represented the corresponding proliferative rate *ρ*_*T*_(fig. S9(B)) for the distinct groups with positive versus negative survival benefit finding a strong size effect (Cohen’s d = -1.31 [-6.0, -0.3]). We later used the value of *ρ*_*T*_as a binary clasification for the two outcome groups and obtained an AUC = 0.96 in the resulting ROC curve, confirming that there is association of low proliferation rates to be benefited from protracted IR, while high proliferative gliomas show a worse outcome under protracted IR.

Altogether, these results indicate that the biology represented by our model supports IR fractionation for medium and low-aggressive tumors, though providing only a minimal survival gain. However, this does not imply that other biological factors may not become more relevant in the context of fractionation, potentially leading to survival benefits when applied.

### 3.6. Tumor cell killing by macrophages

Having established that radiation therapy is necessary for glioma control but not curative, we investigated the macrophage’s ability to counter tumor growth at different stages and under varying IR protocols. Specifically, we examined the tumor cell killing rate exerted by anti-tumor macrophages *A* on tumor cells *T*. This rate includes direct phagocytosis by macrophages and indirect mechanisms, such as collaboration with T cells [10]. Using eq. (S10) from SI, we calculated the percentage of tumor cells *T* eliminated by antitumor macrophages *A* at different time points: before the application of IR (“Pre-IR”), at the midpoint of the simulated time for untreated cases (“*t*_1*/*2,no IR_”), and at the midpoint of the simulated time of the same digital twins undergoing radiation therapy (“*t*_1*/*2,IR_”). The time point *t*_1*/*2,no IR_ was consistently earlier than *t*_1*/*2,IR_ for every mice digital twin, and was used it to analyze both mice undergoing IR and untreated mice, allowing for comparisons at equivalent time points.

As shown in fig. 5, the overall tumor cell killing rates predicted by our model are low, consistently below 0.6 %. Natural tumor progression seemed to foster an immunosuppressive TME for cases with higher tumor cell killing, with a decrease from “Pre-IR” to the midpoint of untreated tumor growth (“No IR, *t*_1*/*2,no IR_” in fig. 5(A)). However, the effect size was inconclusive with the CI of Cohen’s d containing the value zero, probably due to the slight increases in tumor cell killing detected for the low value cases. Under IR application (Cohen’s d = 0.3 [0.1, 0.4]), macrophage-mediated tumor cell killing at the midpoint of untreated tumor growth (“Standard IR, *t*_1*/*2,no IR_”) was lower than at the midpoint of treated tumor growth (“Standard IR, *t*_1*/*2,IR_”).

**Figure 5.**
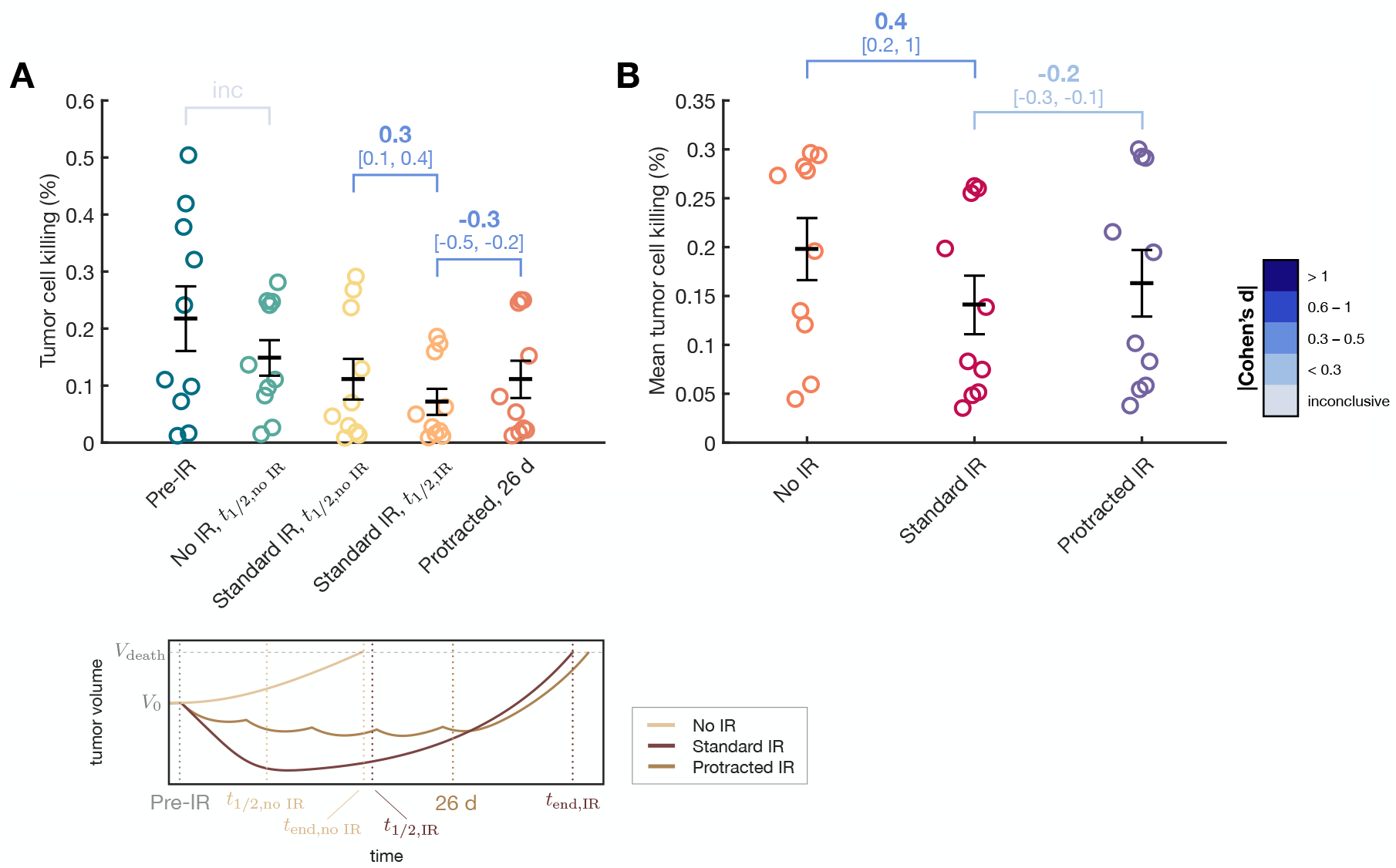
Tumor cell killing effect of anti-tumor macrophages in the in silico model. (A) Percentage of tumor cells *T* eliminated by anti-tumor macrophages *A* at the different time points illustrated in the lower figure: before IR treatment (Pre-IR), midpoint of untreated mice time (*t*_1*/*2,no IR_, half of survival time with no treatment *t*_end,no IR_), midpoint of mice treated with IR (*t*_1*/*2,IR_, half of survival time under IR *t*_end,IR_), and after protracted IR treatment with 6-day interval between fractions (26 d). (B) Time-averaged tumor cell killing rate for untreated mice digital twins (No IR), mice undergoing 5 fractions of daily radiation therapy (Standard IR), and mice undergoing 5 fractions of radiation therapy at 6-day intervals (Protracted IR). Graphs show means *±* SEM. Effect size is reported using Cohen’s d, along with its confidence interval shown in square brackets.

Overall, IR promoted an immunosuppressive TME, resulting in reduced tumor cell killing rates. The effect had a medium effect size (Cohen’s d = 0.4 [0.2, 1]), as tumor cell killing was already very low in every case. Across the simulation duration, the time-averaged tumor cell killing (fig. 5(B)) was higher in untreated tumors (mean 0.2 %) compared to those undergoing standard daily IR (mean 0.14 %). Protracted IR with a 6-day interval very slightly improved mean tumor cell killing (Cohen’s d = -0.2 [-0.3, -0.1]), although the absolute values remained within the low ranges described in this section.

### 3.7. Tackling macrophage phagocytic activity improves survival

Given that the overall tumor-killing activity of macrophages in our model is very low, we wondered whether targeting the phagocytic activity of macrophages could have therapeutic value in gliomas, and what effect our model would predict for such an intervention. Glioma cells are known to upregulate the expression of the surface protein CD47, which acts as a ‘don’t eat me’ signal, inhibiting phagocytosis by binding to its receptor SIRP*α* on MDM and MG [51]. Research has shown that this signaling axis can be blocked, for example, with anti-CD47 monoclonal antibodies [52]. In a GBM xenograft model, anti-CD47 treatment increased tumor cell phagocytosis from 2.7 % to 13.3 %, representing a 5-fold increase [53].

To study the influence of anti-CD47 treatment on the digital twins of the experimental mice used in this study, we simulated its effect by modifying the tumor killing parameter *µ* in the ODE model. To err on the side of caution, we applied a multiplier of 3 to this parameter, which is smaller than the values reported in the previously cited mouse model. We ran simulations for the digital twins undergoing IR treatment alone and for those undergoing both IR and anti-CD47 treatment. The corresponding Kaplan-Meier curves (fig. 6) show that combining IR with anti-CD47 would increase survival compared to IR alone, with a median survival difference of 10 days. The benefit appears to be more pronounced in medium and long survivors (inset of fig. 6) and smaller in IR poor responders. It should be noted that although the difference between curves is not significant (log-rank p-value = 0.11), the difference has to be taken into account, as we are comparing the same digital twins undergoing two treatment modalities rather than performing a trial with randomized individuals.

**Figure 6.**
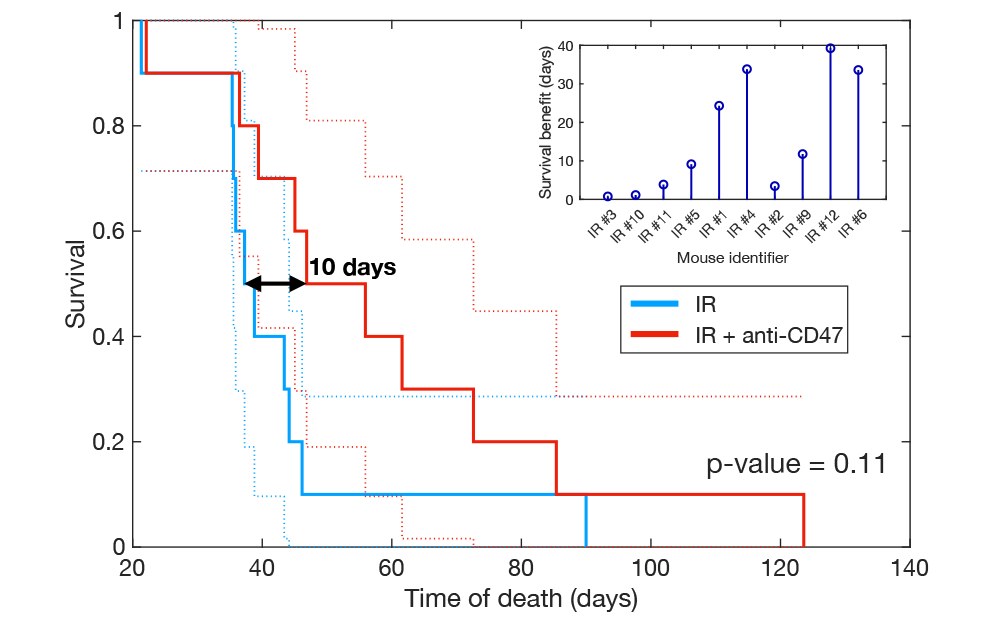
Survival benefit of treatment with anti-CD47. Enhancing macrophage phagocytic activity with anti-CD47 treatment, in combination with IR, provides a survival benefit compared to IR alone. The inset shows the individual survival gains for each of the mice digital twins, ordered by survival time with IR alone from lowest to highest.

Treatment with anti-CD47 alone in silico provided some benefit compared to untreated mice (fig. S7(A)); however, its effect on reducing tumor volume was minimal (fig. S7(B)), and much smaller than IR effect (Cohen’s d = -0.9 [-3.1, -0.5]). Therapies targeting phagocytic activity should be combined with debulking treatments, such as IR. Mathematically, this is because IR directly targets the tumor populations, while anti-CD47 affects their derivatives, thus limiting the overall therapeutic effect.

We found no evidence of correlation between survival benefit from IR + anti-CD47 treatment and the log-transformed proliferation rate *ρ*_*T*_(*R*^2^ = 0.11, F-test p-value = 0.34; see fig. S9(C)). Similarly, *ρ*_*T*_was not a good classifier of mice benefiting from the treatment (Cohen’s d = -0.2 [-2.1, 0.8] and AUC = 0.6; see fig. S9(D)).

Overall, these results demonstrate that targeting the ability of TAMs to recognize and kill tumor cells holds therapeutic value that can complement radiation therapy.

### 3.8. Myeloid cell immunotherapies targeting MDM influx enhance survival

After checking that myeloid cell immunotherapies targeted at the capability of macrophages to phagocytize tumor cells are effective against gliomas in silico, we asked whether other forms of immunotherapy aimed to TAMs can enhance survival. From the previous results presented here we knew that IR alters the populations in the TME and partly causes an increase in TAMs due to an elevated external flux of MDMs. We wondered whether blocking the IR-induced increase in the flux of infiltrating macrophages could have therapeutic value. Leukocyte adhesion and trafficking into tissue during inflammation are mediated by very late antigen-4 (VLA-4), among other factors, which forms a heterodimer of CD49d and integrin beta-1 (CD29) [54, 55]. When leukocytes are activated by chemotactic agents or other stimuli, VLA-4 adheres to its primary ligands, VCAM-1 and fibronectin [56]. Consequently, it has been proposed that neutralizing CD49d, the integrin alpha-4 subunit, limits immune cell entry into the brain [45, 57]. This can be achieved using monoclonal antibodies that neutralize CD49d and disrupt leukocyte adhesion and migration (*α*-CD49d) [58].

To simulate the effect of *α*-CD49d, we focused on the effect of IR on the parameter *ϕ* which controls the external influx of MDMs. As explained in section 2, this influx varies after IR by a factor inverse to 15 the survival fraction of each mouse. With *α*-CD49d, we assumed this factor is uniformly reduced across all mice, thereby limiting the inflammatory effect of IR.

We simulated the application of IR + *α*-CD49d in all mouse digital twins and studied the changes in the distribution of macrophages between the anti-tumor *A* and the pro-tumor *P* populations (fig. 7(A)). The simulations showed that *α*-CD49d increased the proportion of anti-tumor macrophages by 14 percentage points at the expense of pro-tumor macrophages (Cohen’s d = 1.0 [0.7, 1.6]). Additionally, we evaluated the effect on glioma cell proliferation and found that blocking the influx of external macrophages post-IR reduced proliferation by 39 % at 21 days after treatment initiation (Cohen’s d = 0.9 [0.3, 2.4]; see fig. 7(B)). However, at the time of recurrence, the proliferation rates were similar between digital twins treated with and without *α*-CD49d (Cohen’s d = 0.1 [0.05, 0.2]).

**Figure 7.**
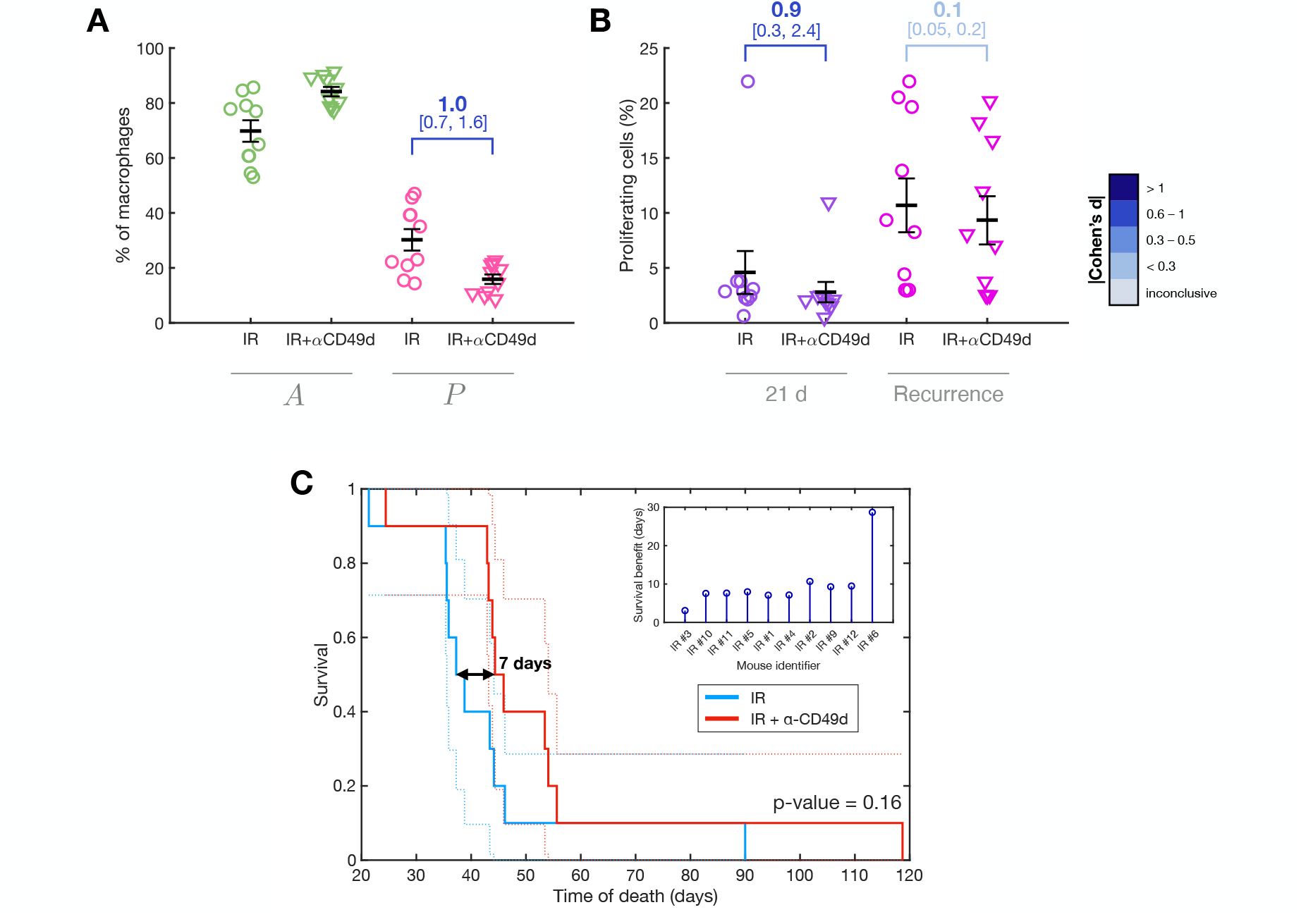
Benefit of treatment with *α*-CD49d. Blocking the influx of infiltrating MDMs induced by IR enhanced the efficacy of radiotherapy. (A) Proportion of anti-tumor and pro-tumor macrophages among total macrophages in mouse digital twins treated with IR or with IR + *α*-CD49d. (B) Percentage of proliferating glioma cells at two time points: 21 days after treatment initiation and at tumor recurrence. Graphs (A) and (B) show means *±* SEM. Effect size is reported using Cohen’s d, along with its confidence interval shown in square brackets. (C) Survival differences between the two groups; the inset shows individual survival gains due to *α*-CD49d.

These changes impacted the survival of groups treated with IR + *α*-CD49d compared to IR alone. Every simulated mouse in the IR + *α*-CD49d group showed a consistent life extension, with an average gain of 7 days (fig. 7(C)). The survival gain was relatively uniform across the group, except for the longest-surviving mouse, which experienced a survival extension three times greater than the average (30 days, see fig. 7(C)-inset). Similar to the effect of anti-CD47, the survival difference was not statistically significant (log-rank p-value = 0.16); however, since the subjects compared are identical copies, the observed survival gain can be considered meaningful.

We also analyzed the correlation between the survival benefit from IR + *α*-CD49d treatment and the log-transformed proliferation rate *ρ*_*T*_finding no linear relationship (*R*^2^ = 0.05, F-test p-value = 0.55; see fig. S9(E)). There was also no difference in *ρ*_*T*_between groups with low and high benefit from IR + *α*-CD49d treatment (Cohen’s d = -0.1 [-6.6, 0.9], and AUC = 0.52; see fig. S9(F)), evidencing that proliferation rate is not able to discriminate subjects that would benefit from the treatment.

These results demonstrate that blocking external MDM influx is a viable therapeutic strategy to enhance the efficacy of IR in gliomas.

### 3.9. CSF-1R inhibition as a therapeutic strategy

Beyond the myeloid cell-targeted immunotherapies described earlier, other interventions targeting fundamental macrophage pathways have been proposed. CSF-1 is the primary growth factor regulating macrophage differentiation, survival, proliferation, and activation [59]. It binds to the receptor CSF-1R, expressed on macrophage surfaces, activating downstream signaling pathways. CSF-1 plays a critical role in TAM activation and their transition to an pro-tumor phenotype [60]. Given the central role of TAMs in tumor biology, targeting either CSF-1 or its receptor has been suggested as a therapeutic strategy for gliomas [45, 61] and other tumors [62].

To conclude our study on TAM-targeted therapies for glioma treatment, we evaluated the application of a CSF-1R inhibitor. Unlike the previously considered interventions, which focused on very specific aspects of TAM biology, CSF-1R inhibition has broad implications and a more complex mechanism of action. This complexity increases the modeling challenge, as multiple terms are influenced by its effects. Based on the findings reported in the literature, we considered three main effects of CSF-1R inhibition:

1. Increase in the phagocytic activity of anti-tumor macrophages [61]. In the same way as in the anti CD47, we do this by modifying the value of the tumor killing parameter *µ* upon CSF-1R inhibition treatment.
2. Deployment of the TAMs influx caused by IR [45]. Following the modeling we used for the effect of *α*CD49d, we modify the increase in *ϕ* after IR exposure.
3. Inversion of the M1-M2 polarization [45, 63]. CSF-1R inhibition has been found to revert the polarization of TAMs and, as a consequence, reduce tumor proliferation. Upon treatment with CSF-1R inhibitors, we inverted the *A* → *P* transition through the change of sign and magnitude in *λ*.

We simulated the effects of a CSF-1R inhibitor on PDG-Ink4a/ArfKO mice digital twins, both as a standalone treatment and in combination with IR. The CSF-1R inhibitor was administered over 12 days, starting at the same day that the first IR fraction for the combination therapy. Biomarker effects were assessed at the end of the treatment period of 12 days. Tumor cell killing by anti-tumor macrophages increased by 80 % in both treatment modalities compared to IR alone (Cohen’s d = -0.5 [-1.1, -0.3] for CSF-1R inhibitor and Cohen’s d = -0.4 [-0.5, -0.2] for IR + CSF-1R; see fig. 8(A)), driven by an increase in anti-tumor macrophages and their enhanced ability to eliminate tumor cells. Glioma cell proliferation was reduced by half with the CSF-1R inhibitor alone (Cohen’s d = 1.4 [0.6, 3.7]) and decreased nearly fivefold when combined with IR (Cohen’s d = 2.9 [1.0, 9.1]; see fig. 8(B)). This is due to the reduced influence of pro-tumor macrophages implicated in the proliferation feedback loop. Additionally, macrophage populations shifted toward an anti-tumor phenotype, with a 10-percentage-point increase in *A* with the inhibitor alone ((Cohen’s d = 0.5 [0.2, 1.6])) and a 32-point increase when used as an IR adjuvant (Cohen’s d = 1.6 [0.8, 3.6]; see fig. 8(C)).

**Figure 8.**
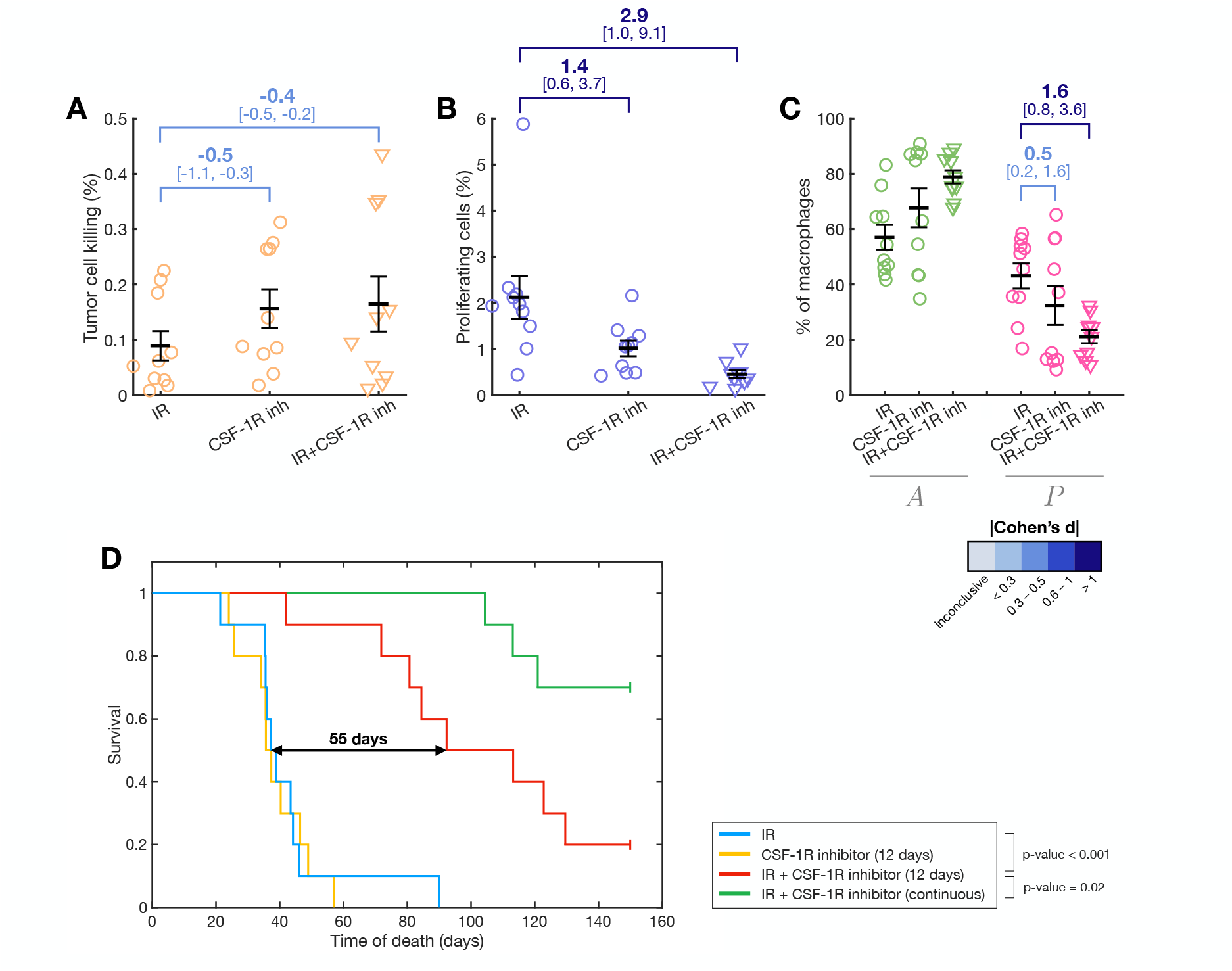
Effect of CSF-1R inhibition on virtual mice. Simulation results for the following treatments: 5 daily fractions of IR alone, 12 days of CSF-1R inhibitor alone, and their combination, measured 12 days after treatment initiation. (A) Tumor cell killing mediated by anti-tumor macrophages. (B) Glioma cell proliferation rates. (C) Macrophage distribution between anti-tumor (*A*) and pro-tumor (*P*) populations. Graphs (A-C) show means *±* SEM. Effect size is reported using Cohen’s d, along with its confidence interval shown in square brackets. (D) Survival outcomes from in silico clinical trials of these treatments.

Combining the CSF-1R inhibitor with IR significantly improved survival in virtual mice, demonstrating their synergistic effects (fig. 8(D)). However, the CSF-1R inhibitor alone did not enhance survival compared to IR alone. To assess the impact of prolonged treatment, we simulated a protocol with continuous CSF 1R inhibitor administration. This approach yielded a marked increase in life expectancy (fig. 8(D), green Kaplan-Meier curve). Model predictions were compared with experimental data from ref. [45], reproduced in fig. S8, for completeness. While the model qualitatively aligned with trends observed in real mice, it over estimated survival for the 12-day CSF-1R + 5-day IR protocol and underestimated survival with continuous CSF-1R application. Despite these discrepancies, the model effectively captured the therapeutic potential of this myeloid cell-targeted immunotherapy, paving the way for future refinements and applications.

To finalize, we checked whether individuals with higher or lower proliferation rate could differentially benefit from 12-day CSF-1R + 5-day IR administration. We found a significant linear correlation between the survival benefit and the log-transformed tumor proliferation rate (*R*^2^ = 0.76, F-test p-value=0.005; see fig. S9(G)). Using the proliferation rate *ρ*_*T*_as a binary classifier for the high benefit and low benefit subgroups separated by the survival benefit median yielded and AUC = 0.96, with a effect size of -0.8 [-2.0, -0.2] as quantified by Cohen’s d (fig. S9(H)). This indicates that the success of the combined treatment of IR with adjuvant CSF-1R is affected by the value of the proliferation rate (lower values being associated to a better outcome) in our in silico model.

## 4. Discussion

Macroscopic glioma growth is driven by microscopic interactions between various cell types within the TME. TAMs, in particular, represent the most abundant immune population in the TME. TAMs play a pivotal role in glioma progression by promoting tumor cell proliferation and motility, supporting angiogenesis, and inducing immune suppression, among other effects [9, 64, 65]. Macrophages are therefore a cornerstone of glioma biology. Furthermore, they are involved in the response to multiple treatments, including chemotherapy, radiotherapy, and immunotherapy [11, 19]. Therapeutic interventions targeting the specific traits of TAMs are attracting significant interest and research, emerging as a rapidly growing field [4, 64, 66]. Myeloid cells in general, and TAMs in particular, are anticipated to be central to the next wave of cancer immunotherapy [1]. In this context, mathematical and computational models are crucial for systematizing biological knowledge, performing simulations, optimizing therapeutic applications, and ultimately enabling personalized medicine [67].

In this work, we developed a minimal model of glioma growth that integrates the key interactions between tumor cells and TAMs, accounting for both MG and MDM. Importantly, the model incorporates the positive feedback loop sustaining tumor growth through M2-like macrophages [12]. The model successfully fits available data on the volumetric dynamics of experimental gliomas in PDG-Ink4a/ArfKO mice, with the proliferation rate of tumor cells as the sole fitting parameter for untreated cases, and the addition of the survival fraction for treated cases. Moreover, the model qualitatively reproduces trends in various TAM biomarkers. To maintain the simplicity of the model, we excluded other potentially important cell populations, such as monocytes or T cells; however, the framework presented here can easily be extended to include these and other populations.

To explore the effects of different treatment modalities, we used longitudinal data from experimental mice to create digital twins—virtual representations that replicate tumor progression in the reference individuals [68]. Using these mouse digital twins, we compared the outcomes of various treatment variations by simulating the progression of the same virtual subject under different treatments or schemes. This approach allows for comparisons within the same individual, unlike in real-life clinical trials, where comparisons must be made across randomized groups of individuals, each with their own biological variability. As a result, interventions that produce differences within the same digital twin can be considered relevant, even if statistical tests designed for random samples fail to detect those differences. Nevertheless, it is important to note that the complete information about tumor growth, response and survival in each mouse was already known at the time the digital twin was created. Thus, this study does not involve the creation of real-time digital twins capable of predicting the future behavior of partially observed clinical cases, which still presents challenges [69].

Notably, we incorporated the effect of IR into our framework for three main reasons. Firstly, radiation therapy is a critical component of current glioma treatments and can not be overlooked [70, 71]. Secondly, IR induces significant changes in the tumor immunological state, particularly regarding myeloid cell infiltration, making its inclusion essential when these cells play a key role [14]. Thirdly, to date myeloid-cell-targeted immunotherapies are more likely to be administered in combination with debulking options rather than as standalone treatments [72]; from a mathematical perspective, this is because IR modeling affects absolute population levels, while immunotherapy effects are reflected in specific terms of the population derivatives. Our results of IR simulations showed an improvement in the survival of in silico mice by 18 days, consistent with experimental findings. We also explored the application of protracted therapy with varying time intervals, finding a slight benefit in cases with a median prognosis, though not in poor survivors. This benefit, however, would likely be undetectable in a randomized clinical trial. This does not necessarily imply that protracted schemes are ineffective, but rather that the biology captured in our model does not suggest significant alterations under such regimens. Finally, while it would have been interesting to study the effects of different IR doses, this was challenging due to the lack of data needed to infer behavior. A classical approach, such as using the linear quadratic (LQ) model to evaluate dose-dependent differences in survival, could have been applied [73]. However, the impact of varying doses on TAM populations remains impossible to assess without further experimental research.

We used our model to simulate the effects of novel immunotherapies targeting TAMs, which show promise as part of the therapeutic arsenal against gliomas. Specifically, we focused on three types of therapies: those targeting the phagocytic ability of TAMs in the TME (anti-CD47), the infiltration of MDMs in response to IR-mediated inflammation (*α*CD49d), and inhibitors of CSF-1R, which disrupt the interaction between TAMs and the key cytokine CSF-1, affecting multiple aspects of TAM behavior. Our findings indicate that these therapies exhibit promising therapeutic effects when combined with IR in digital copies of PDG Ink4a/ArfKO mice. When experimental data were available, the model results showed good agreement with the experimental observations. Based on these results, we believe that the current version of the model will be valuable in the development and pre-clinical assessment of these therapies, as it provides a framework for simulating and analyzing their effects and exploring variations. The model enables the development of in silico clinical trials and serves as a tool for therapy optimization. Furthermore, it can be easily extended to incorporate additional therapies.

We expect this line of work to eventually translate into clinical applications. Patient data can be incorporated to extend and refine the current model. For example, positron emission tomography (PET) is currently being investigated as a tool to image macrophage presence [74], and other data sources, such as histopathological samples, can provide insights into the state of the TME [75–77]. However, using patient data comes with limitations; for instance, untreated or clearly sub-optimal cases are unlikely to be available as baseline references. In previous work, we have used imaging data to inform mathematical modeling, which has helped propose modifications to therapeutic protocols and optimize treatments [47]. This approach also has the advantage of identifying global patterns that might be missed when focusing solely on individual cases [78]. Our ultimate goal is to move toward patient-specific personalization, investigating how individual subjects would respond to treatments and how their specific situations can be improved. Although there are inherent differences between the behavior of human gliomas and their animal model counterparts [65], we believe that this animal-based initial model can serve as a valuable reference for future applications in patients.

A reasonable assumption would be to consider that tumor cells could also grow in vitro the absence of macrophages. Yet, in many cases, up to 50% of cells in the glioma microenvironment are macrophages [79]. In this context, modeling tumor growth in a total absence of macrophages becomes an abstraction mostly valid under laboratory conditions, while our work focuses on scenarios where tumor growth is strongly impacted by interactions with macrophages (see SI 1). Moreover, and because these interactions are mostly mediated by diffusive chemical secretions and do not require explicit cell contact, we model them proportional to cellular abundances and not their effective densities, which is the classical mean-field population approach followed by most models (see SI 1 and e.g. [80–82]). A high cellular density, however, is still detrimental to tumor progression due to the saturation shape of Gompertzian growth.

Despite its achievements, our model has limitations. First, it simplifies the glioma TME considerably by excluding potentially relevant cell populations and their diversity [1]. For instance, we neglected monocytes, which differentiate into MDMs, assuming rapid transformation with minimal functional impact, and lacking data or markers to quantify this population in the used dataset. Similarly, we omitted T cells and other lymphocytes, which are known to play roles in the TME, even in brain tumors [83]. While we partially accounted for macrophage-mediated T cell action as part of the anti-tumor macrophages action, the explicit role of lymphocytes was overlooked. When brought together, all this evidence points towards the need of understanding tumors as high-dimensional complex systems, in which far more than 2 or 3 cell types interact [84, 85]. To do so, we need to incorporate mathematical framework for species-rich ecological communities, which reveal that non-trivial dynamics can emerge solely from the interactions between such a large number of cancer and non-cancer cells [85].

Moreover, there is a high uncertainty in the values of the parameters due to the lack of literature data on their individualized values, particularly for the fixed parameters. In principle, an automated optimization method could be used to fit both the volumetric time evolution and the bulk biomarker data, for instance by using the average values of the distributions for validation. However, this approach also poses challenges such as defining a suitable objective function or keeping accuracy both in the volumetric evolution and in each of the available biomarkers given the limited information available. For this reason, we adopted a simplified heuristic method which ensures a good agreement between average experimental and simulated data while maintaining simplicity (see section 2.5).

Another limitation is that our model does not account for spatial dynamics. The ODE-based framework used here simplifies tumor-immune system interactions to a non-spatial context. Extending these rules to an agent-based model (ABM) of whole tumor growth could provide insights into the spatial heterogeneity of tumor growth and immune responses [86, 87]. Complementarily, the study of tumor-macrophage interactions at the microscale with detailed ABMs can also contribute to the elucidation of spatiotemporal dynamics at the lower scale [88]. Finally, we treated TAMs as existing in binary M1-like or M2-like phenotypic states. A more sophisticated approach would model the spectrum of TAM phenotypes and incorporate additional dimensions of their behavior, reflecting the complexity of their roles in the TME [81, 89, 90]. Cross regulatory interactions between MG and MDM in the TME are also biologically plausible, for instance through reciprocal cytokine signaling, and might be a relevant addition to future models.

In conclusion, we have developed a glioma growth model that captures key cancer-macrophage inter actions in the TME. The model incorporates the effects of radiation therapy and simulates the impact of promising myeloid cell-targeted immunotherapies. Using published data from experiments using PDG Ink4a/ArfKO mice, we parametrized the model to ensure biological relevance. This framework can support the development of next-generation immunotherapies by enabling simulation-based exploration and computational optimization. This work provides a foundation that can be further developed for studies tailored to individual patients, opening avenues for personalized treatment strategies.

## Acknowledgements

The authors thank Leila Akkari for her availability and in-depth explanations of TAM biology. The authors thank Alejandro Bertolet for discussion about glioma response to radiation. The authors thank Mara de Martino for her support in understanding the role of MDMs in tumors. The authors thank Víctor M. Pérez-García for discussion about the mathematical model. The authors thank Eric Holland and Frank Szulzewsky for their detailed explanation of gliomagenesis in animal models. The authors thank Beatriz Ocaña-Tienda for discussions on statistical methods and graphical representation.

## Code availability

The source codes to produce the results and analyses presented in this manuscript are available on the following Github repository: https://github.com/jesusjbosque/myeloid-immunotherapies.git

## References

[1] S. Goswami, S. Anandhan, D. Raychaudhuri, P. Sharma, Myeloid cell-targeted therapies for solid tumours, Nature Reviews Immunology 23 (2) (2023) 106–120.

[2] D. G. DeNardo, B. Ruffell, Macrophages as regulators of tumour immunity and immunotherapy, Nature Reviews Immunology 19 (6) (2019) 369–382.

[3] D. J. Kloosterman, L. Akkari, Macrophages at the interface of the co-evolving cancer ecosystem, Cell 186 (8) (2023) 1627–1651.

[4] S. T. Barry, D. I. Gabrilovich, O. J. Sansom, A. D. Campbell, J. P. Morton, Therapeutic targeting of tumour myeloid cells, Nature Reviews Cancer 23 (4) (2023) 216–237.

[5] F. De Sanctis, A. Adamo, S. Can‘e, S. Ugel, Targeting tumour-reprogrammed myeloid cells: the new battleground in cancer immunotherapy, Seminars in Immunopathology 45 (2) (2023) 163–186.

[6] A. Buonfiglioli, D. Hambardzumyan, Macrophages and microglia: the cerberus of glioblastoma, Acta neuropathologica communications 9 (2021) 1–21.

[7] D. F. Quail, J. A. Joyce, The microenvironmental landscape of brain tumors, Cancer cell 31 (3) (2017) 326–341. doi:10.1016/j.ccell.2017.02.009.

[8] B. M. Andersen, C. Faust Akl, M. A. Wheeler, E. A. Chiocca, D. A. Reardon, F. J. Quintana, Glial and myeloid heterogeneity in the brain tumour microenvironment, Nature Reviews Cancer 21 (12) (2021) 786–802.

[9] D. Hambardzumyan, D. H. Gutmann, H. Kettenmann, The role of microglia and macrophages in glioma maintenance and progression, Nature neuroscience 19 (1) (2016) 20–27.

[10] P. Schol, M. J. van Elsas, J. Middelburg, M. K. N. Twilhaar, T. van Hall, T. C. van der Sluis, S. H. van der Burg, Myeloid effector cells in cancer, Cancer Cell 42 (12) (2024) 1997–2014.

[11] C. Engblom, C. Pfirschke, M. J. Pittet, The role of myeloid cells in cancer therapies, Nature Reviews Cancer 16 (7) (2016) 447–462.

[12] K. V. Myers, K. J. Pienta, S. R. Amend, Cancer cells and m2 macrophages: Cooperative invasive ecosystem engineers, Cancer Control 27 (1) (2020) 1073274820911058.

[13] J. Raizer, A. Parsa, et al., Current understanding and treatment of gliomas, Vol. 163, Springer, 2015.

[14] R. A. Bekker, S. Kim, S. Pilon-Thomas, H. Enderling, Mathematical modeling of radiotherapy and its impact on tumor interactions with the immune system, Neoplasia 28 (2022) 100796.

[15] H. E. Barker, J. T. Paget, A. A. Khan, K. J. Harrington, The tumour microenvironment after radiotherapy: mechanisms of resistance and recurrence, Nature Reviews Cancer 15 (7) (2015) 409–425.

[16] L. Bejarano, M. J. Jordőo, J. A. Joyce, Therapeutic targeting of the tumor microenvironment, Cancer discovery 11 (4) (2021) 933–959.

[17] K. Mortezaee, M. Najafi, Immune system in cancer radiotherapy: Resistance mechanisms and therapy perspectives, Critical reviews in oncology/hematology 157 (2021) 103180.

[18] C. Beach, D. MacLean, D. Majorova, J. N. Arnold, M. M. Olcina, The effects of radiation therapy on the macrophage response in cancer, Frontiers in oncology 12 (2022) 1020606.

[19] S. A. Lasser, F. G. Ozbay Kurt, I. Arkhypov, J. Utikal, V. Umansky, Myeloid-derived suppressor cells in cancer and cancer therapy, Nature Reviews Clinical Oncology 21 (2) (2024) 147–164.

[20] D. Barbolosi, J. Ciccolini, B. Lacarelle, F. Barlési, N. André, Computational oncology—mathematical modelling of drug regimens for precision medicine, Nature reviews Clinical oncology 13 (4) (2016) 242–254.

[21] J. D. Butner, P. Dogra, C. Chung, R. Pasqualini, W. Arap, J. Lowengrub, V. Cristini, Z. Wang, Mathematical modeling of cancer immunotherapy for personalized clinical translation, Nature computational science 2 (12) (2022) 785–796.

[22] R. J. De Boer, P. Hogeweg, H. Dullens, R. A. De Weger, W. Den Otter, Macrophage T lymphocyte interactions in the anti-tumor immune response: a mathematical model., Journal of Immunology (Baltimore, Md.: 1950) 134 (4) (1985) 2748–2758.

[23] M. R. Owen, J. A. Sherratt, Pattern formation and spatiotemporal irregularity in a model for macrophage–tumour interactions, Journal of Theoretical Biology 189 (1) (1997) 63–80.

[24] M. R. Owen, J. A. Sherratt, Modelling the macrophage invasion of tumours: Effects on growth and composition, Mathematical Medicine and Biology: A Journal of the IMA 15 (2) (1998) 165–185.

[25] H. Byrne, S. Cox, K.C.E., Macrophage-tumour interactions: In vivo dynamics, Discrete and Continuous Dynamical Systems - B 4 (1) (2004) 81–98.

[26] H. Knútsdóttir, E. Pálsson, L. Edelstein-Keshet, Mathematical model of macrophage-facilitated breast cancer cells invasion, Journal of Theoretical Biology 357 (2014) 184–199.

[27] N. Y. den Breems, R. Eftimie, The re-polarisation of M2 and M1 macrophages and its role on cancer outcomes, Journal of Theoretical Biology 390 (2016) 23–39.

[28] K.-A. Norton, K. Jin, A. S. Popel, Modeling triple-negative breast cancer heterogeneity: Effects of stromal macrophages, fibroblasts and tumor vasculature, Journal of Theoretical Biology 452 (2018) 56–68.

[29] G. Mahlbacher, L. T. Curtis, J. Lowengrub, H. B. Frieboes, Mathematical modeling of tumor-associated macrophage interactions with the cancer microenvironment, Journal for immunotherapy of cancer 6 (2018) 1–17.

[30] X. Li, M. K. Jolly, J. T. George, K. J. Pienta, H. Levine, Computational modeling of the crosstalk between macrophage polarization and tumor cell plasticity in the tumor microenvironment, Frontiers in oncology 9 (2019) 10.

[31] F. Leonard, L. T. Curtis, A. R. Hamed, C. Zhang, E. Chau, D. Sieving, B. Godin, H. B. Frieboes, Nonlinear response to cancer nanotherapy due to macrophage interactions revealed by mathematical modeling and evaluated in a murine model via CRISPR-modulated macrophage polarization, Cancer Immunology, Immunotherapy 69 (2020) 731–744.

[32] R. Eftimie, L. Gibelli, A kinetic theory approach for modelling tumour and macrophages heterogeneity and plasticity during cancer progression, Mathematical Models and Methods in Applied Sciences 30 (04) (2020) 659–683.

[33] L. Bartha, R. Eftimie, Mathematical investigation into the role of macrophage heterogeneity on the temporal and spatiotemporal dynamics of non-small cell lung cancers, Journal of Theoretical Biology 549 (2022) 111207.

[34] S. Mayer, T. Milo, A. Isaacson, C. Halperin, S. Miyara, Y. Stein, C. Lior, M. Pevsner-Fischer, E. Tzahor, A. Mayo, et al., The tumor microenvironment shows a hierarchy of cell-cell interactions dominated by fibroblasts, Nature Communications 14 (1) (2023) 5810.

[35] P. Shojaee, F. Mornata, A. Deutsch, M. Locati, H. Hatzikirou, The impact of tumor associated macrophages on tumor biology under the lens of mathematical modelling: A review, Frontiers in Immunology 13 (2022) 1050067.

[36] P. Tracqui, G. Cruywagen, D. Woodward, G. Bartoo, J. Murray, E. Alvord Jr, A mathematical model of glioma growth: the effect of chemotherapy on spatio-temporal growth, Cell proliferation 28 (1) (1995) 17–31.

[37] K. R. Swanson, C. Bridge, J. Murray, E. C. Alvord Jr, Virtual and real brain tumors: using mathematical modeling to quantify glioma growth and invasion, Journal of the neurological sciences 216 (1) (2003) 1–10.

[38] H. Hatzikirou, A. Deutsch, C. Schaller, M. Simon, K. Swanson, Mathematical modelling of glioblastoma tumour development: a review, Mathematical Models and Methods in Applied Sciences 15 (11) (2005) 1779–1794.

[39] V.M. Pérez-García, G. F. Calvo, J. Belmonte-Beitia, D. Diego, L. Pérez-Romasanta, Bright solitary waves in malignant gliomas, Physical Review E—Statistical, Nonlinear, and Soft Matter Physics 84 (2) (2011) 021921.

[40] J. Alfonso, K. Talkenberger, M. Seifert, B. Klink, A. Hawkins-Daarud, K. Swanson, H. Hatzikirou, A. Deutsch, The biology and mathematical modelling of glioma invasion: a review, Journal of the Royal Society Interface 14 (136) (2017) 20170490.

[41] M. Conte, Y. Dzierma, S. Knobe, C. Surulescu, Mathematical modeling of glioma invasion and therapy approaches via kinetic theory of active particles, Mathematical Models and Methods in Applied Sciences 33 (05) (2023) 1009–1051.

[42] M. Elitaş, S. Zeinali, Modeling and simulation of EGF-CSF-1 pathway to investigate glioma-macrophage interaction in brain tumors, International Journal of Cancer Studies & Research (IJCR): Special Issue On” Advances in Brain Cancer Research” 5 (2016) 1–8.

[43] Y. Kim, H. Jeon, H. Othmer, The role of the tumor microenvironment in glioblastoma: A mathematical model, IEEE Transactions on Biomedical Engineering 64 (3) (2016) 519–527.

[44] P. Shojaee, E. Weinholtz, N. S. Schaadt, F. Feuerhake, H. Hatzikirou, Biopsy location and tumor-associated macrophages in predicting malignant glioma recurrence using an in-silico model, npj Systems Biology and Applications 11 (1) (2025) 3.

[45] L. Akkari, R. L. Bowman, J. Tessier, F. Klemm, S. M. Handgraaf, M. de Groot, D. F. Quail, L. Tillard, J. Gadiot, J. T. Huse, et al., Dynamic changes in glioma macrophage populations after radiotherapy reveal csf-1r inhibition as a strategy to overcome resistance, Science translational medicine 12 (552) (2020) eaaw7843.

[46] K. R. Swanson, R. C. Rostomily, E. Alvord, A mathematical modelling tool for predicting survival of individual patients following resection of glioblastoma: a proof of principle, British journal of cancer 98 (1) (2008) 113–119.

[47] T. Delobel, L.E. Ayala-Hernández, J. J. Bosque, J. Pérez-Beteta, S. Chulián, M. García-Ferrer, P. Piñero, P. Schucht, M. Murek, V.M. Pérez-García, Overcoming chemotherapy resistance in low-grade gliomas: A computational approach, PLoS computational biology 19 (11) (2023) e1011208.

[48] B. Segura-Collar, J. Jiménez-Sánchez, R. Gargini, M. Dragoj, J.M. Sepúlveda-Sánchez, M. Pešić, M. A. Ramírez, L.E. Ayala-Hernández, P. Sánchez-Gómez, V.M. Pérez-García, On optimal temozolomide scheduling for slowly growing glioblastomas, Neuro-Oncology Advances 4 (1) (2022) vdac155.

[49] A. Henares-Molina, S. Benzekry, P. C. Lara, M. García-Rojo, V.M. Pérez-García, A. Martínez-González, Non-standard radiotherapy fractionations delay the time to malignant transformation of low-grade gliomas, PLoS One 12 (6) (2017) e0178552.

[50] S. C. Brüningk, J. Peacock, C. J. Whelan, R. Brady-Nicholls, H.-H. M. Yu, S. Sahebjam, H. Enderling, Intermittent radiotherapy as alternative treatment for recurrent high grade glioma: A modeling study based on longitudinal tumor measurements, Scientific reports 11 (1) (2021) 20219.

[51] M. J. Pittet, O. Michielin, D. Migliorini, Clinical relevance of tumour-associated macrophages, Nature reviews Clinical oncology 19 (6) (2022) 402–421.

[52] S. B. Willingham, J.-P. Volkmer, A. J. Gentles, D. Sahoo, P. Dalerba, S. S. Mitra, J. Wang, H. Contreras-Trujillo, R. Martin, J. D. Cohen, et al., The cd47-signal regulatory protein alpha (sirpa) interaction is a therapeutic target for human solid tumors, Proceedings of the National Academy of Sciences 109 (17) (2012) 6662–6667.

[53] G. Hutter, J. Theruvath, C. M. Graef, M. Zhang, M. K. Schoen, E. M. Manz, M. L. Bennett, A. Olson, T. D. Azad, R. Sinha, et al., Microglia are effector cells of cd47-sirpα antiphagocytic axis disruption against glioblastoma, Proceedings of the National Academy of Sciences 116 (3) (2019) 997–1006.

[54] K. A. Pacheco, M. Tarkowski, J. Klemm, L. J. Rosenwasser, Cd49d expression and function on allergen-stimulated t cells from blood and airway, American journal of respiratory cell and molecular biology 18 (2) (1998) 286–293.

[55] L. A. Haile, J. Gamrekelashvili, M. P. Manns, F. Korangy, T. F. Greten, Cd49d is a new marker for distinct myeloid-derived suppressor cell subpopulations in mice, The Journal of Immunology 185 (1) (2010) 203–210.

[56] B. Chan, M. Elices, E. Murphy, M. Hemler, Adhesion to vascular cell adhesion molecule 1 and fibronectin. comparison of alpha 4 beta 1 (vla-4) and alpha 4 beta 7 on the human b cell line jy., Journal of Biological Chemistry 267 (12) (1992) 8366–8370.

[57] K. J. Myers, D. R. Witchell, M. J. Graham, S. Koo, M. Butler, T. P. Condon, Antisense oligonucleotide blockade of alpha 4 integrin prevents and reverses clinical symptoms in murine experimental autoimmune encephalomyelitis, Journal of neuroimmunology 160 (1-2) (2005) 12–24.

[58] K.-C. Lin, A. C. Castro, Very late antigen 4 (vla4) antagonists as anti-inflammatory agents, Current Opinion in Chemical Biology 2 (4) (1998) 453–457.

[59] A. Sehgal, K. M. Irvine, D. A. Hume, Functions of macrophage colony-stimulating factor (csf1) in development, homeostasis, and tissue repair, in: Seminars in immunology, Vol. 54, Elsevier, 2021, p. 101509.

[60] E. R. Stanley, V. Chitu, Csf-1 receptor signaling in myeloid cells, Cold Spring Harbor perspectives in biology 6 (6) (2014) a021857.

[61] S. M. Pyonteck, L. Akkari, A. J. Schuhmacher, R. L. Bowman, L. Sevenich, D. F. Quail, O. C. Olson, M. L. Quick, J. T. Huse, V. Teijeiro, et al., CSF-1R inhibition alters macrophage polarization and blocks glioma progression, Nature medicine 19 (10) (2013) 1264–1272. doi:10.1038/nm.3337.

[62] M. A. Cannarile, M. Weisser, W. Jacob, A.-M. Jegg, C. H. Ries, D. Rüttinger, Colony-stimulating factor 1 receptor (csf1r) inhibitors in cancer therapy, Journal for immunotherapy of cancer 5 (1) (2017) 53.

[63] J. H. Stafford, T. Hirai, L. Deng, S. B. Chernikova, K. Urata, B. L. West, J. M. Brown, Colony stimulating factor 1 receptor inhibition delays recurrence of glioblastoma after radiation by altering myeloid cell recruitment and polarization, Neuro-oncology 18 (6) (2016) 797–806.

[64] A. Mantovani, F. Marchesi, A. Malesci, L. Laghi, P. Allavena, Tumour-associated macrophages as treatment targets in oncology, Nature reviews Clinical oncology 14 (7) (2017) 399–416.

[65] F. Klemm, R. R. Maas, R. L. Bowman, M. Kornete, K. Soukup, S. Nassiri, J.-P. Brouland, C. A. Iacobuzio-Donahue, C. Brennan, V. Tabar, Interrogation of the microenvironmental landscape in brain tumors reveals disease-specific alterations of immune cells, Cell 181 (7) (2020) 1643–1660. doi:10.1016/j.cell.2020.05.007.

[66] A. Mantovani, P. Allavena, F. Marchesi, C. Garlanda, Macrophages as tools and targets in cancer therapy, Nature Reviews Drug Discovery 21 (11) (2022) 799–820.

[67] R. Eftimie, A. Mavrodin, S. P. Bordas, From digital control to digital twins in medicine: A brief review and future perspectives, Advances in Applied Mechanics 56 (2023) 323–368.

[68] C. Wu, G. Lorenzo, D. A. Hormuth, E. A. Lima, K. P. Slavkova, J. C. DiCarlo, J. Virostko, C. M. Phillips, D. Patt, C. Chung, et al., Integrating mechanism-based modeling with biomedical imaging to build practical digital twins for clinical oncology, Biophysics reviews 3 (2) (2022).

[69] T. Hernandez-Boussard, P. Macklin, E. J. Greenspan, A. L. Gryshuk, E. Stahlberg, T. Syeda-Mahmood, I. Shmulevich, Digital twins for predictive oncology will be a paradigm shift for precision cancer care, Nature medicine 27 (12) (2021) 2065–2066.

[70] A. R. Cabrera, J. P. Kirkpatrick, J. B. Fiveash, H. A. Shih, E. J. Koay, S. Lutz, J. Petit, S. T. Chao, P. D. Brown, M. Vogelbaum, et al., Radiation therapy for glioblastoma: executive summary of an american society for radiation oncology evidence-based clinical practice guideline, Practical Radiation Oncology 6 (4) (2016) 217–225.

[71] E. P. Sulman, N. Ismaila, T. S. Armstrong, C. Tsien, T. T. Batchelor, T. Cloughesy, E. Galanis, M. Gilbert, V. Gondi, M. Lovely, et al., Radiation therapy for glioblastoma: American society of clinical oncology clinical practice guideline endorsement of the american society for radiation oncology guideline, Journal of Clinical Oncology 35 (3) (2017) 361–369.

[72] A. S. Ding, D. Routkevitch, C. Jackson, M. Lim, Targeting myeloid cells in combination treatments for glioma and other tumors, Frontiers in immunology 10 (2019) 1715.

[73] W. Sung, T. S. Hong, M. C. Poznansky, H. Paganetti, C. Grassberger, Mathematical modeling to simulate the effect of adding radiation therapy to immunotherapy and application to hepatocellular carcinoma, International Journal of Radiation Oncology* Biology* Physics 112 (4) (2022) 1055–1062.

[74] Y. C. Toner, G. Prévot, M. M. van Leent, J. Munitz, R. Oosterwijk, A. V. D. Verschuur, Y. van Elsas, V. Peric, R. J. Maas, A. Ranzenigo, et al., Macrophage pet imaging in mouse models of cardiovascular disease and cancer with an apolipoprotein-inspired radiotracer, npj Imaging 2 (1) (2024) 12.

[75] J. A. Bull, E. J. Mulholland, J. W. Moore, J. J. Bosque, B. J. Stolz, J. Boen, H. R. Eggington, H. L. Belnoue-Davis, H. Jones, C. D. Gatenbee, et al., Integrating diverse statistical methods to analyse stage-discriminatory cell interactions in colorectal neoplasia, bioRxiv (2024) 2024–06.

[76] J. A. Bull, J. W. Moore, E. J. Mulholland, S. J. Leedham, H. M. Byrne, Muspan: A toolbox for multiscale spatial analysis, bioRxiv (2024) 2024–12.

[77] Y. Azimzade, M. H. Haugen, V. N. Kristensen, A. Frigessi, A. Kohn-Luque, Integrated multiomics analysis unveils how macrophages drive immune suppression in breast tumors and affect clinical outcomes, bioRxiv (2024) 2024–11.

[78] J. J. Bosque, G. F. Calvo, D. Molina-García, J. Pérez-Beteta, A. M. G. Vicente, V.M. Pérez-García, Metabolic activity grows in human cancers pushed by phenotypic variability, Iscience 26 (3) (2023).

[79] F. Khan, L. Pang, M. Dunterman, M. S. Lesniak, A. B. Heimberger, P. Chen, et al., Macrophages and microglia in glioblastoma: heterogeneity, plasticity, and therapy, The Journal of clinical investigation 133 (1) (2023).

[80] V. A. Kuznetsov, I. A. Makalkin, M. A. Taylor, A. S. Perelson, Nonlinear dynamics of immunogenic tumors: parameter estimation and global bifurcation analysis, Bulletin of mathematical biology 56 (2) (1994) 295–321.

[81] R. Eftimie, C. Barelle, Mathematical investigation of innate immune responses to lung cancer: The role of macrophages with mixed phenotypes, Journal of Theoretical Biology 524 (2021) 110739.

[82] S. Sabir, O. Leòn-Triana, S. Serrano, R. Barrio, V.M. Pérez-García, Mathematical model of car t-cell therapy for a b-cell lymphoma lymph node, Bulletin of Mathematical Biology 87 (3) (2025) 1–33.

[83] M. B. Watowich, M. R. Gilbert, M. Larion, T cell exhaustion in malignant gliomas, Trends in cancer 9 (4) (2023) 270–292.

[84] B. P. Kotler, J. S. Brown, Cancer community ecology, Cancer Control 27 (1) (2020) 1073274820951776.

[85] G. Aguadé-Gorgorió, A. R. Anderson, R. Solé, Modeling tumors as complex ecosystems, Iscience 27 (9) (2024).

[86] J. Jiménez-Sánchez, Á. Martínez-Rubio, A. Popov, J. Pérez-Beteta, Y. Azimzade, D. Molina-García, J. Belmonte-Beitia, G. F. Calvo, V.M. Pérez-García, A mesoscopic simulator to uncover heterogeneity and evolutionary dynamics in tumors, PLoS computational biology 17 (2) (2021) e1008266.

[87] L. V. Kunz, J. J. Bosque, M. Nikmaneshi, I. Chamseddine, L. L. Munn, J. Schuemann, H. Paganetti, A. Bertolet, Amber: A modular model for tumor growth, vasculature and radiation response, Bulletin of Mathematical Biology 86 (12) (2024) 139.

[88] J. A. Bull, H. M. Byrne, Quantification of spatial and phenotypic heterogeneity in an agent-based model of tumour-macrophage interactions, PLOS Computational Biology 19 (3) (2023) e1010994.

[89] R. M. Ransohoff, A polarizing question: do m1 and m2 microglia exist?, Nature neuroscience 19 (8) (2016) 987–991. doi:10.1038/nn.4338.

[90] C. Yunna, H. Mengru, W. Lei, C. Weidong, Macrophage m1/m2 polarization, European journal of pharmacology 877 (2020) 173090. doi:10.1016/j.ejphar.2020.173090.

