## Supplementary Material for "Mathematical model of tumor-macrophage dynamics in glioma to advance myeloid-targeted therapies"

July 2025

#### SI1 Assumptions of the mathematical model

Our mathematical model is based on several assumptions, most of which are supported by the available literature. Below, we list the assumptions made.

- The immune landscape of gliomas is dominated by myeloid cells and, in particular tumor-associated macrophages (TAMs) composed of microglia (MG) and monocyte-derived macrophages (MDMs) [1, 2]. We simplified our modeling framework to consider only tumor cells, and TAMs.
- We consider that both MG and MDM, besides having distinct origins, can be considered together in a common population of macrophages within the TME. This is because it is not entirely clear whether both cell types have different functions in the brain TME [3] and the adaptations of both types of cells make it difficult to ascribe them unique functions [4, 5].
- Some macrophages, and in particular MG, are able to exert cytotoxic and phagocytic effects and could eliminate tumor cells [4]. Induced changes in TAM gene expression with phenotypes showing enhanced phagocytosis of glioma cells resulted in a more pronounced effect of tumor reduction compared with TAM depletion [6, 3] which suggests that macrophages in the TME have the ability to combat the tumor under certain circumstances. Therefore, we splitted the TAM population in an anti-tumor (sometimes called M1-like [7]) subpopulation $A$ and a pro-tumor subpopulation $P$ (M2-like) [8].
- There is an influx of external MDMs from the blood circulation [9] that we model by the flux parameter $\phi$.
- The main way in which gliomas recruit MDMs is by the secretion of periostin produced by glioma stem cells (but not so much by differentiated glioma cells), as MDM density correlates with periostin levels in human GBMs and silencing it in glioma stem cells reduces their density and inhibits tumor growth [8]. Therefore, we assume that the influx of MDMs increases with the number of tumor cells.
- The subtype of macrophages recruited from the blood is M2-like, promoting tumor growth [8].
- Both MDM and MG subpopulations proliferate within the tumor microenvironment (TME). They do so in response to the presence of tumor cells [4].
- Pro-tumor TAM subpopulations are known to facilitate and promote tumor proliferation [10, 11, 4, 9, 8], therefore, we assume here that tumor cells proliferate at a faster rate as the number of pro-tumor TAMs grows.
- A first assumption would be to separate tumor growth into two components, $(\rho_i + \rho_T P)$, related to intrinsic and macrophage-mediated growth rates. Yet, because gliomas are highly populated by macrophages [12], we do not have relevant *in vivo* information about $\rho_i$. In fact, such intrinsic growth

rate is likely relevant only under laboratory conditions where cancer cells can be grown isolated from other cellular interactions with different populations from the tumor microenvironment. To focus on those scenarios where macrophages are abundant and key to glioma dynamics [2], we assume here that $\rho_i << \rho_T P$, and focus on studying the dynamics where $\rho_i$ is negligible. A natural next step to our model could be to infer the dynamics of tumor cells inside and outside of their microenvironment to be able to estimate the differences between intrinsic and macrophage-mediated growth.

- Most of the macrophage–tumor interactions in gliomas described above are mediated by non-local, paracrine signaling rather than direct cell-cell contact. Glioma cells secrete cytokines and chemokines that recruit and polarize macrophages, while macrophages release factors that promote tumor proliferation and invasion [8, 4, 9]. As such, we assume that interaction terms depend on total cell numbers rather than local concentrations [13, 14, 15]. To account for spatial and density constraints, we include a saturation term, $\log(K/V)$, which reduces the effect of interactions (and hence growth) at high cell densities. This approach captures both the diffuse non-local nature of the interactions and the crowding limitations in the tumor microenvironment.
- Ionizing radiation (IR) of gliomas causes a change towards the immunosuppressive state [16]. High-dose and low-dose irradiation increase infiltration and polarization of macrophages to suppressive phenotypes [17, 18]. In the model, IR causes an increase in the influx of pro-tumor macrophages proportional to the killing effect of IR on tumor cells.
- TAMs are highly resistant to ionizing radiation [18]. In the model, the survival fraction of TAMs is bigger than the survival fraction of tumor cells.

### SI2 Individual tracking of MG and MDM

The ordinary differential equations (ODE) model presented in the main text, eqs. (1) to (3), classifies tumor and immune cells by their functional role within the TME. However, it is convenient to keep track of independent numbers of MG and MDM to enable comparisons with biological data. With this purpose in mind, here we separate the population of pro-tumor macrophages into the population that comes from transformed MG ($P_1$) and those composed of externally infiltrated MDM ($P_2$). Thus, eq. (3) can be separated into the contribution of

$$\frac{\mathrm{d}P_1}{\mathrm{d}t} = \rho_P P_1 \cdot T \cdot \log\left(\frac{K}{V}\right) + \lambda A \cdot T \frac{K_\lambda}{K_\lambda + T} - \frac{1}{\tau_M} P_1, \tag{S1}$$

$$\frac{\mathrm{d}P_2}{\mathrm{d}t} = \rho_P P_2 \cdot T \cdot \log\left(\frac{K}{V}\right) + \phi \cdot T - \frac{1}{\tau_M} P_2, \tag{S2}$$

with

$$P = P_1 + P_2. \tag{S3}$$

### SI3 Data from experiments in mice

We contrast and parametrize our model with empirical data from experiments on PDG-Ink4a/ArfKO immunocompetent mice previously published in the literature [2]. The generation of the animal model has been previously described in ref. [6]. Briefly, genetically engineered mouse models (GEMMs) were injected with chicken fibroblasts DF-1 engineered to express the RCAS-PDGFB and RCAS-Cre vectors. Glial cells in the GEMMs are infected with the RCAS virus, which targets neural progenitor cells, leading to the over-expression of PDGFB in the GEMMs and subsequent glioma formation *de novo* facilitated by the deletion of Ink4a/Arf suppressor genes. Therefore, gliomas in this model arise spontaneously without external injection of tumor cells and the mice are immunocompetent, allowing the study of tumor-immune interactions.

##### SI3.1 Tumor volume measurement

Tumor volumes were sequentially measured with T2-weighted magnetic resonance imaging (MRI) scans. After tumor formation, mice underwent fractionated radiation therapy at a dose of 2 Gy per day for 5 days. Four mice were used as control and imaged at 3 time points. Twelve mice, numbered 1 to 12 here, underwent radiotherapy and were imaged a variable number of times (between 2 and 7) until the mouse either died from tumor progression or had to be euthanized due to neurological symptoms. For these mice, the total tumor volume as measured per MRI was available. For mice #7 and #8 only two volumetric time points were available and they were excluded from our study.

##### SI3.2 Measurement of biomarkers

Additionally, there is available information on specific biomarkers on mice following the same treatment protocol. The three main ones were the percentage of Iba1$^+$ cells, corresponding to the macrophages present in the tumor, the percentage of Ki67$^+$ cells within the Iba1$^-$ fraction, marking proliferation of tumor cells, and percentage of CC3$^+$ cells, indicating the proportion of apoptotic cells. These data were available for four time points for Ki67$^+$ and Iba1$^+$: control, after 5 days of IR (5 d), 21 days after the initiation of IR (21 d), and at tumor recurrence (IR Rec). For CC3$^+$, only the control and 5 d were available. Tumor recurrence was defined as the tumor reaching a threshold of 90 mm$^3$ or presenting with neurological disorders that led to the mouse being humanely euthanized.

Moreover, to distinguish between resident MG and infiltrating MDM, the authors of ref. [2] measured the percentages of these two groups within the total macrophages (Iba1$^+$ cells) by flow cytometry quantification of CD49d$^-$ cells (MG) and CD49d$^+$ (MDMs). These data were available for the control, at the end of 5 days of fractionated IR, and at recurrence.

The data on tumor volume growth were individualized for each mouse, allowing for detailed analysis at the individual level. In contrast, biomarker data were provided in bulk, without information on their temporal progression for individual mice.

#### SI4 Calculation of markers from the model

The data and metrics emerging from the experiments described above need to be mapped to the variables in our model. To do so, we define the variables presented below.

- **Macrophages:** We counted the percentage of macrophages, which in the experiments from ref. [2] were identified by cells expressing Iba1, simply by calculating the percentage of the total populations made up by the macrophage populations:

  $$\text{macrophages}(t) = \frac{P(t) + A(t) + M_D(t)}{N(t)} \times 100, \tag{S4}$$

  where

  $$N(t) = T(t) + P(t) + A(t) + T_D(t) + M_D(t) \tag{S5}$$

  is the total number of cells in the system.

  Regarding their distribution into MG and MDM, we took advantage of the internal separation in $P_1$ (made up entirely of transformed MG) and $P_2$ (accounting for externally infiltrating MDM) used within the simulation executions (see section SI2) to quantify the distribution of these two populations within the total number of macrophages:

  $$\text{MG}(t) = \frac{P_1(t) + A(t)}{P(t) + A(t)} \times 100, \tag{S6}$$

  $$\text{MDM}(t) = \frac{P_2(t)}{P(t) + A(t)} \times 100. \tag{S7}$$

- **Proliferating cells:** The proliferation of glioma cells was measured in ref. [2] as the percentage of the total cells that were Ki67$^+$ Iba1$^-$, that is, glioma cells undergoing mitosis. To translate this measure we calculated the new glioma cells generated by mitosis by integrating the proliferation term from eq. (1) from the main text (i.e., $\rho_T PT \log(K/V)$) over the mitosis duration $\Delta t_{\text{prol}}$. This number of new cells was then normalized by the initial number of cells at the beginning of that time period ($N(t - \Delta t_{\text{prol}})$):

$$\text{proliferation}(t) = \frac{\int_{t-\Delta t_{\text{prol}}}^{t} \rho_T PT \log\left(\frac{K}{V}\right)\, \mathrm{d}\tau}{N(t-\Delta t_{\text{prol}})} \times 100. \tag{S8}$$

- **Apoptotic cells:** In the experiments, apoptotic cells were identified as the percentage of cells expressing CC3. In our model, there are two sources of dying cells, which we attribute to apoptosis in both cases. One is the decay of damaged cells due to treatment and the other is the killing of tumor cells by anti-tumor macrophages. The first is simply calculated by the number of damaged cells that leave the system during the characteristic time for death $\Delta t_{\text{apop}}$, while the second is given by integrating the disappearing tumor cells over that time due to the killing term:

$$\text{apoptosis}(t) = \frac{(T_D(t-\Delta t_{\text{apop}}) - T_D(t)) + (M_D(t-\Delta t_{\text{apop}}) - M_D(t)) + \int_{t-\Delta t_{\text{apop}}}^{t} \mu TA\, \mathrm{d}\tau}{N(t-\Delta t_{\text{apop}})} \times 100. \tag{S9}$$

- **Tumor cell killing:** Similarly, we can track the percentage of tumor cells eliminated by anti-tumor macrophages $A$ at a given time. This killing encompasses both the direct phagocytosis of tumor cells by macrophages and other macrophage-mediated mechanisms, such as cooperation with T cells [7], which are not explicitly modeled here. To calculate this rate, we integrate the corresponding term and divide it by the initial number of tumor cells, as follows:

$$\text{killing}(t) = \frac{\int_{t-\Delta t_{\text{apop}}}^{t} \mu TA\, \mathrm{d}\tau}{T(t-\Delta t_{\text{apop}})} \times 100. \tag{S10}$$

Throughout the article, we have used unfilled markers to illustrate the previous metrics calculated from in silico simulations.

### SI5 Basic properties of the mathematical model

We state the following proposition.

**Proposition 1** *For any positive initial data $(T(0), P(0), A(0))$ and all parameters of the initial value problem given by eqs. (1) to (3) of the main text being positive, there exists $h > 0$ such that solutions for $T(t), P(t), A(t)$ exist for $t \in [-h, h]$, are unique and positive.*

**Proof 1** *Local existence and uniqueness follow straightforwardly from the Picard-Lindelöf theorem, since the right-hand side of system (1) to (3) is continuous and locally Lipschitz. With regard to the positivity of the solutions, eq. (1) can be written in an integral form*

$$T(t) = T(0)e^{\int_0^t \rho_T T(s)P(s)\log(K/V(s)) - \mu T(s)A(s)ds} > 0$$

*if $T(0) > 0$.*

*Regarding $P(t)$ and $A(t)$, we reason in contradiction: Let us assume that there exists a $t_1 > 0$ value such that $P(t_1) = 0$ and $\frac{dP}{dt}(t_1) < 0$, being $t_1$ the first time it happens.*

*Then, for this case*

$$\frac{dP}{dt}(t_1) = \lambda A(t_1)\frac{K_\lambda T(t_1)}{K_\lambda + T(t_1)} + \phi T(t_1) < 0$$

*and, by the positivity of $T(t)$, we infer $A(t_1) < 0$. Since $A(0) > 0$ and $A$ is a continuous function, there exists a $t_0 > 0$, $t_0 < t_1$ such that $A(t_0) = 0$ and $dA(t_0)/dt < 0$.*

*Then*

$$\frac{dA}{dt}(t_0) = \xi \frac{dT(t_0)}{dt} < 0$$

*but*

$$\frac{dT}{dt}(t_0) = \rho_T T(t_0) P(t_0) \ln \frac{K}{V(t_0)} > 0$$

*since $P(t) > 0$ for $t \in (0, t_1)$. So, we conclude the proof.*

■

Now, we calculate the equilibrium points of system eqs. (1) to (3) and its stability. Solving the equations

$$\rho_T T \cdot P \cdot \log\left(\frac{K}{V}\right) - \mu T A = 0, \tag{S11}$$

$$\rho_P P \cdot T \cdot \log\left(\frac{K}{V}\right) + \lambda A \cdot T \frac{K_\lambda}{K_\lambda + T} + \phi \cdot T - \frac{1}{\tau_M} P = 0, \tag{S12}$$

$$\rho_A A \cdot T \cdot \log\left(\frac{K}{V}\right) - \lambda A \cdot T \frac{K_\lambda}{K_\lambda + T} + \xi \frac{\mathrm{d}T}{\mathrm{d}t} - \frac{1}{\tau_M} A = 0, \tag{S13}$$

we obtain the following equilibrium points:

$$M = (T_1, P_1, A_1) = (0, 0, 0), \tag{S14}$$

$$N = (T_2, P_2, A_2) = \left(\frac{K}{v_T + v_M \phi \tau_M}, \frac{K}{\frac{v_T}{\phi \tau_M} + v_M}, 0\right) \tag{S15}$$

$$Q = (T_3, P_3, A_3) = (T^*, P^*, A^*) \tag{S16}$$

where the last equilibrium points satisfy the equation:

$$\frac{\mu}{\rho_T}(\rho_P - \rho_A) A^* + \lambda A^* \frac{K_\lambda}{K_\lambda + T^*} + \phi = -\lambda P^* \frac{K_\lambda}{K_\lambda + T^*}$$

Since $\rho_P > \rho_A$, the only values that satisfy this relationship are negative values for some of the components $T^*$, $P^*$, $A^*$ in the equation. Since they have no biological meaning, they will not be considered here.

To analyze the stability of these points, we calculate the Jacobian matrix of eqs. (1) to (3):

$$J = \begin{pmatrix} \rho_T P \log\left(\frac{K}{V}\right) - \rho_T T P \left(\frac{v_T}{V}\right) - \mu A & \rho_T T \log\left(\frac{K}{V}\right) - \rho_T T P \left(\frac{v_M}{V}\right) & -\rho_T T P \left(\frac{v_M}{V}\right) - \mu T \\ \rho_P P \log\left(\frac{K}{V}\right) - \rho_P P T \left(\frac{v_T}{V}\right) + \lambda A \frac{K_\lambda}{K_\lambda + T} + \phi & \rho_P T \log\left(\frac{K}{V}\right) - \rho_P P T \left(\frac{v_M}{V}\right) - \frac{1}{\tau_M} & -\rho_P P T \left(\frac{v_M}{V}\right) + \lambda T \frac{K_\lambda}{K_\lambda + T} \\ \rho_A A \log\left(\frac{K}{V}\right) - \rho_A A T \left(\frac{v_T}{V}\right) - \lambda A \frac{K_\lambda}{K_\lambda + T} & -\rho_A A T \left(\frac{v_M}{V}\right) & \rho_A T \log\left(\frac{K}{V}\right) - \lambda T \frac{K_\lambda}{K_\lambda + T} - \frac{1}{\tau_M} \end{pmatrix}$$

The jacobian matrix for $M$ is

$$J(M) = \begin{pmatrix} 0 & 0 & 0 \\ \phi & -\frac{1}{\tau_M} & 0 \\ 0 & 0 & -\frac{1}{\tau_M} \end{pmatrix}$$

The eigenvalues are $\lambda_1 = 0$, $\lambda_{2,3} = -\frac{1}{\tau_M}$. Thus, this equilibrium is a non-hyperbolic equilibrium point. To investigate the local stability of this equilibrium point, we take a neighborhood $E(0, 0, 0)$ of the origin, and choose a point in this neighborhood $(\varepsilon_1, \varepsilon_2, 0)$, with $\varepsilon_1, \varepsilon_2$ sufficiently small, and check that for this point, $dT/dt > 0$ and thus, the orbit for which $A = 0$ moves away from the origin, in agreement with the simulations in the main text. Therefore, the origin is an unstable equilibrium point.

The jacobian matrix for $N$ is

$$J(N) = \begin{pmatrix} \frac{-\rho_T \phi \tau_M v_T}{(v_T + v_M \phi \tau_M) K} & \frac{-\rho_T v_M}{v_T + v_M \phi \tau_M} & 0 \\ \frac{-\rho_P \phi \tau_M v_T}{(v_T + v_M \phi \tau_M) K} + \phi & \frac{-\rho_P v_M}{v_T + v_M \phi \tau_M} - \frac{1}{\tau_M} & 0 \\ 0 & 0 & -\frac{1}{\tau_M} \end{pmatrix}$$

The characteristic polynomial for this matrix is

$$p(\lambda) = \left(\lambda + \frac{1}{\tau_M}\right)\left(\lambda^2 + \frac{v_T + \phi\tau_M v_M + \rho_P \tau_M v_M + K\phi\rho_T \tau_M^2 v_T}{\tau_M(v_T + \phi\tau_M v_M)}\lambda + \frac{\phi\rho_T v_M + K\phi\rho_T v_T}{v_T + \phi\tau_M v_M}\right)$$

It is straightforward to check that the roots of the second degree equation are negative, so the three roots of the characteristic polynomial are negative and therefore the equilibrium point

$$N = (T_2, P_2, A_2) = \left(\frac{K}{v_T + v_M\phi\tau_M}, \frac{K}{\frac{v_T}{\phi\tau_M} + v_M}, 0\right)$$

is locally asymptotically stable.

The orbit, which is a solution of the system of differential equations, can be represented in the phase space, as shown in fig. S1 (Left). Note that this orbit is a connection linking the unstable equilibrium point $M$ with the asymptotically stable equilibrium $N$. Figure S1 (Right) shows a zoomed-in view of the equilibrium point $N$, where different trajectories can be seen converging to $N$, along with some vectors indicating the flow of the orbits.

### SI6 Sensitivity analysis

In this section, we present the sensitivity analysis carried out for the model.

#### SI6.1 Non-dimensionalization

In order to carry out a more effective and generalizable sensitivity analysis, the first step is to perform a non-dimensionalization of the system. This aims to simplify the differential equations and highlight the most relevant variables and parameters by removing physical units and expressing the equations into dimensionless quantities. This transformation facilitates the comparison between different scenarios and improves the understanding of the relative influence of each parameter on the dynamic behavior of the system. Furthermore, it allows the identification of characteristic time scales and critical quantities, which can be useful for interpreting the results.

Considering that all parameters are positive, we define the new dimensionless variables as

$$T^* = \frac{T}{C}, \quad P^* = \frac{P}{C}, \quad A^* = \frac{A}{C}, \quad \tau = \mu C t, \qquad \text{where } C = \frac{K}{v_M}.$$

Renaming these for ease of notation, the resulting dimensionless system (Eqs. (1)–(3) of the main text) reads:

$$\begin{cases} \dfrac{dT}{dt} = \rho_T T \cdot P \cdot \log\left(\dfrac{K}{V}\right) - TA, \\ \dfrac{dP}{dt} = \rho_P P \cdot T \cdot \log\left(\dfrac{K}{V}\right) + \lambda A \cdot T \dfrac{K_\lambda}{K_\lambda + T} + \phi \cdot T - \dfrac{1}{\tau_M} P, \\ \dfrac{dA}{dt} = \rho_A A \cdot T \cdot \log\left(\dfrac{K}{V}\right) - \lambda A \cdot T \dfrac{K_\lambda}{K_\lambda + T} + \xi \dfrac{dT}{dt} - \dfrac{1}{\tau_M} A, \end{cases} \tag{S17}$$

where $V = v_T \cdot C \cdot T + v_M \cdot C(P + A)$.

#### SI6.2 Sensitivity analysis procedure

To evaluate the influence of the parameters and initial conditions on the mathematically modeled system, we performed a twofold sensitivity analysis. First, a local analysis was conducted by independently varying each parameter and initial condition across its respective range while keeping the others fixed. The eight parameters analyzed were $(\rho_T, \rho_P, \rho_A, \mu, \lambda, K_\lambda, \phi, \xi)$, and the initial conditions included tumor cells $(T)$, pro-tumor macrophages ($P$), and anti-tumor macrophages ($A$). Each parameter was varied (by varying one parameter at a time) across a range of $\pm 30\%$ from its baseline value, with the limits defined in Table S1. Biologically estimated parameters ($\tau_M, K, v_T, v_M$) were held constant throughout. For the local sensitivity analysis, we used MATLAB's `ode45` numerical solver to solve the system of equations. Parameter intervals were generated by dividing linearly (with $n = 1000$ points) between their respective lower and upper bounds, while initial conditions were varied between the minimum and maximum values observed experimentally. For each simulation, the final state of the cell populations was recorded. Sensitivities were then quantified as the variance of the output solutions corresponding to each parameter and initial condition variation. The relative contribution of each input to the variance was then normalized to yield percentage-based sensitivity scores.

Following this local analysis, a global sensitivity analysis was conducted using Polynomial Chaos Expansions (PCE), implemented in UQLab. The same set of parameters, ranges, and baseline values were used. A metamodel was constructed to approximate the original system efficiently and to estimate Sobol total-order indices with reduced computational cost. This framework considers the full variability range of each parameter and incorporates their probability distributions, providing a more robust and integrated understanding of how each input contributes to output variance, including nonlinearities and interaction effects [19].

#### SI6.3 Results of the sensitivity analysis

The results of the sensitivity analyses are shown in figs. S2 and S3 as bar plots illustrating the contribution of each parameter and the initial conditions to the variance in the populations, $(T, P, A)$ under control and radiotherapy-treated conditions.

Figure S2(a) shows that the parameter $\rho_T$ is the most influential in the three modeled populations of control mice. This suggests that, in the absence of treatment, the system is dominated by the intrinsic aggressiveness of the glioma and its rapid proliferative capacity. Furthermore, the initial conditions $(T_0, P_0, A_0)$ have a limited impact, which could indicate that the glioma and its TME rapidly evolve towards a state where proliferation and tissue invasion become dominant forces. These conclusions remain in the global analysis using Total-Order Sobol Indices (fig. S2(b)) where the influence of $\rho_T$ increases, confirming its overarching role when non-linear interactions are considered. Moreover, the elevated total-order indices for initial conditions imply that feedback between tumor proliferation and immune recruitment amplifies variance, reflecting the dynamic plasticity of the tumor microenvironment when left unperturbed.

In treated mice (fig. S3(a)), the sensitivity to $\rho_T$ increases, highlighting the critical role of controlling tumor proliferation for therapeutic success. Initial conditions $(T_0, P_0, A_0)$ also gain importance, as larger initial tumor cell populations ($T_0$) or pro-tumor macrophages ($P_0$) can hinder treatment efficacy, while higher levels of anti-tumor macrophages ($A_0$) may be beneficial despite their plasticity. The direct effect of $T_0$ on $T$ ($\approx 0.20$) and of $P_0$ on $P$ ($\approx 0.18$) increases, while $A_0$ remains modest ($\approx 0.08$). Treatment exacerbates the instability of the TME, underscoring the need for personalized therapies targeting tumor growth and immune interactions in a dynamic, immunosuppressive context. Interaction parameters $(\lambda, \phi)$ reach moderate influence ($\approx 0.15$)—suggesting some immunomodulatory effect of the treatment—while $(K_\lambda, \varepsilon)$ are around ($\approx 0.10$), indicating limited enhancement of macrophage-mediated tumor cell killing. Both $\rho_T$ and initial conditions converge ($T_0 \approx 0.40, P_0 \approx 0.38$), indicating that under treatment the variance from starting cell counts is almost entirely captured by their interactions, which suggests that precise measurement of tumor burden and macrophage composition prior to radiotherapy is especially important, as their interplay with proliferation and immune-related parameters strongly shapes outcomes.

**Table S1.** Variables and estimated values of the parameters in the mathematical model

| **Description** | **Symbol** | **Value** | **Unit** |
|---|---|---|---|
| Tumor cells | $T$ | Variable | # |
| Anti-tumor macrophages | $A$ | Variable | # |
| Pro-tumor macrophages | $P$ | Variable | # |
| Cell volume | $v_T$, $v_M$ | $5\times10^{-5}$ | $mm^3$ $cell^{-1}$ |
| Tumor proliferation rate | $\rho_T$ | $[3.1\times10^{-7}, 3.2\times10^{-5}]$ | $d^{-1}$ $cell^{-1}$ |
| Survival fraction | $S_T$ | [0.54, 0.87] | - |
| Pro-tumor macrophage proliferation rate | $\rho_P$ | $3.6\times10^{-8}$ | $d^{-1}$ $cell^{-1}$ |
| Anti-tumor macrophage proliferation rate | $\rho_A$ | $1.2\times10^{-8}$ | $d^{-1}$ $cell^{-1}$ |
| Macrophage lifespan | $\tau_M$ | $6 / \ln(2)$ | d |
| Carrying capacity | K | 380 | $mm^3$ |
| Killing rate (control) | $\mu$ | $1.2\times10^{-4}$ | $d^{-1}$ $cell^{-1}$ |
| Killing rate (IR) | $\mu$ | $5\times10^{-7}$ | $d^{-1}$ $cell^{-1}$ |
| Michaelis constant of macrophage transition | $K_\lambda$ | $1.4\times10^{5}$ | cell |
| Macrophage transition rate | $\lambda$ | $2.5\times10^{-7}$ | $d^{-1}$ $cell^{-1}$ |
| MDM flux | $\phi$ | $1.5\times10^{-3}$ | $d^{-1}$ |
| Microglia invasion proportionality | $\xi$ | 0.14 | - |
| Damaged tumor cell lifespan | $\tau_{T_D}$ | 2.2 | d |
| Damaged macrophage lifespan | $\tau_{M_D}$ | 5 | d |

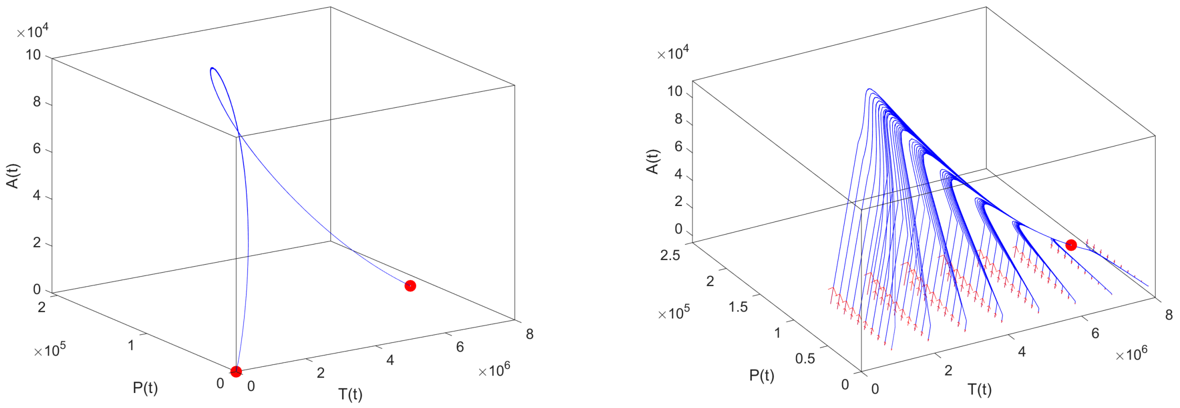


**Figure S1.** (Left) The orbit corresponding to the solution of the system of equations (1), (2), (3) for the initial condition $(T_0, P_0, A_0) = (10^{-3}, 10^{-3}, 10^{-3})$, with parameter values given in table S1. The orbit escapes from the unstable equilibrium $M$ towards the asymptotically stable equilibrium $N$. (Right) Zoom of the equilibrium point $N$, where different trajectories can be seen converging to $N$. The red vectors indicate the direction of the orbit flow.

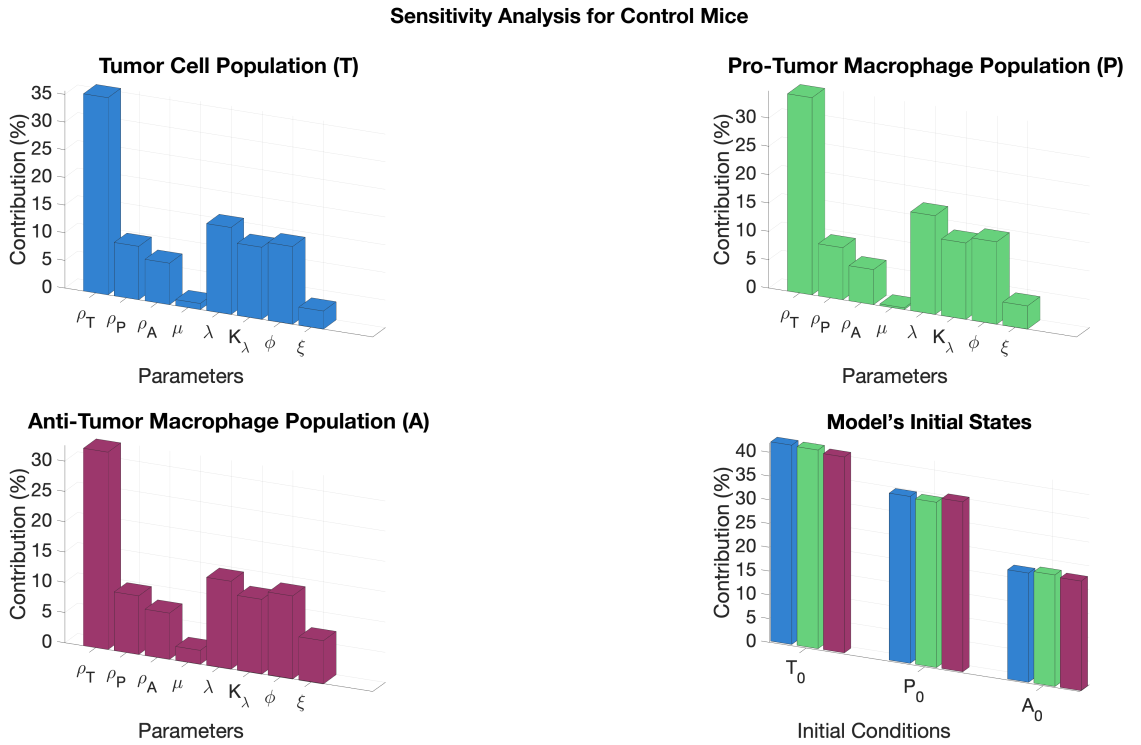


(a) Sensitivity analysis of control mice performed by varying each parameter and initial condition individually across its predefined range. The resulting variances were normalized to reflect the relative influence on model outputs.

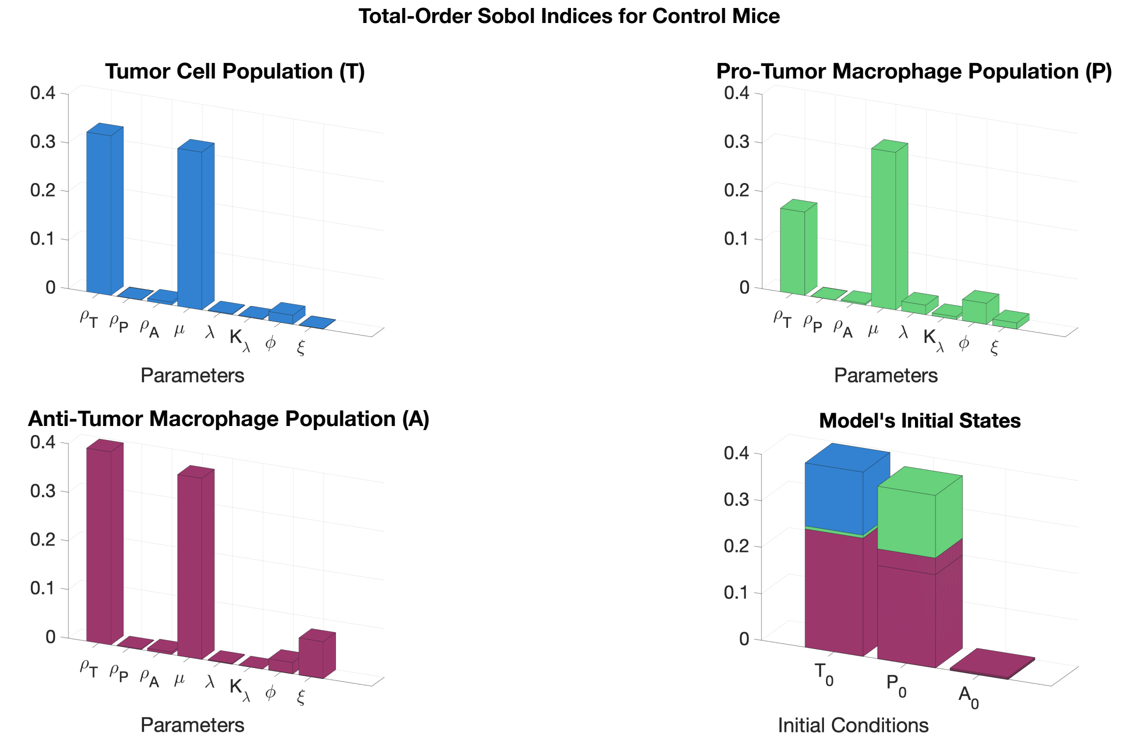


(b) Total-order sensitivity indices for control mice, computed via Polynomial Chaos Expansion (PCE).

**Figure S2.** Sensitivity analysis of control mice using two complementary approaches. Panel (a) shows the variance in model outputs caused by independent variation of each parameter and initial condition. Panel (b) displays total-order Sobol indices obtained via Polynomial Chaos Expansion, capturing the global influence of each parameter including interactions.

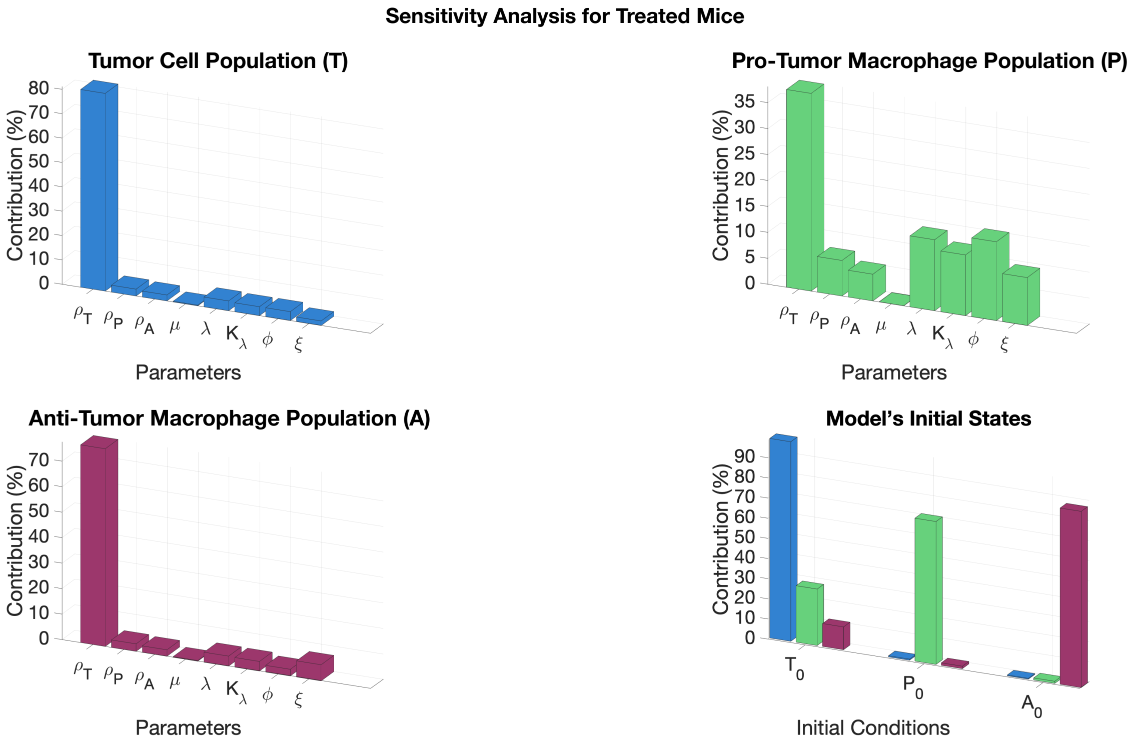


(a) Sensitivity analysis of treated mice performed by varying each parameter and initial condition individually across its predefined range. The resulting variances were normalized to reflect the relative influence on model outputs.

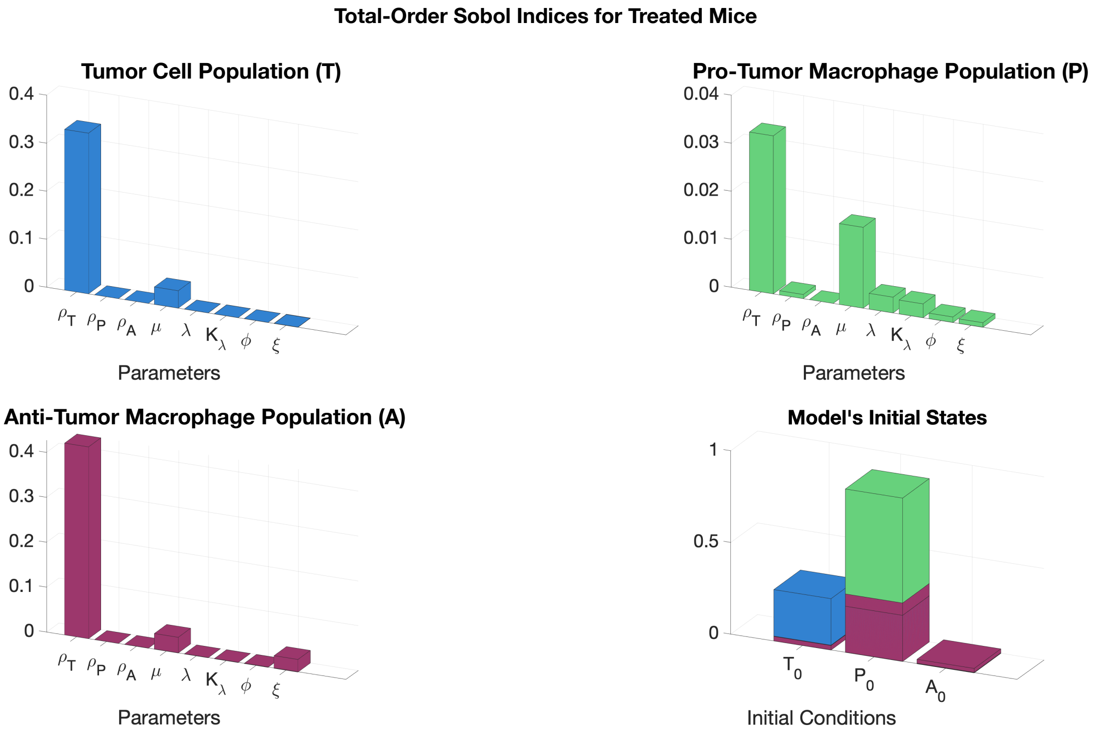


(b) Total-order sensitivity indices for treated mice, computed via Polynomial Chaos Expansion (PCE).

**Figure S3.** Sensitivity analysis of treated mice using two complementary approaches. Panel (a) shows the variance in model outputs caused by independent variation of each parameter and initial condition. Panel (b) displays total-order Sobol indices obtained via Polynomial Chaos Expansion, capturing the global influence of each parameter including interactions.

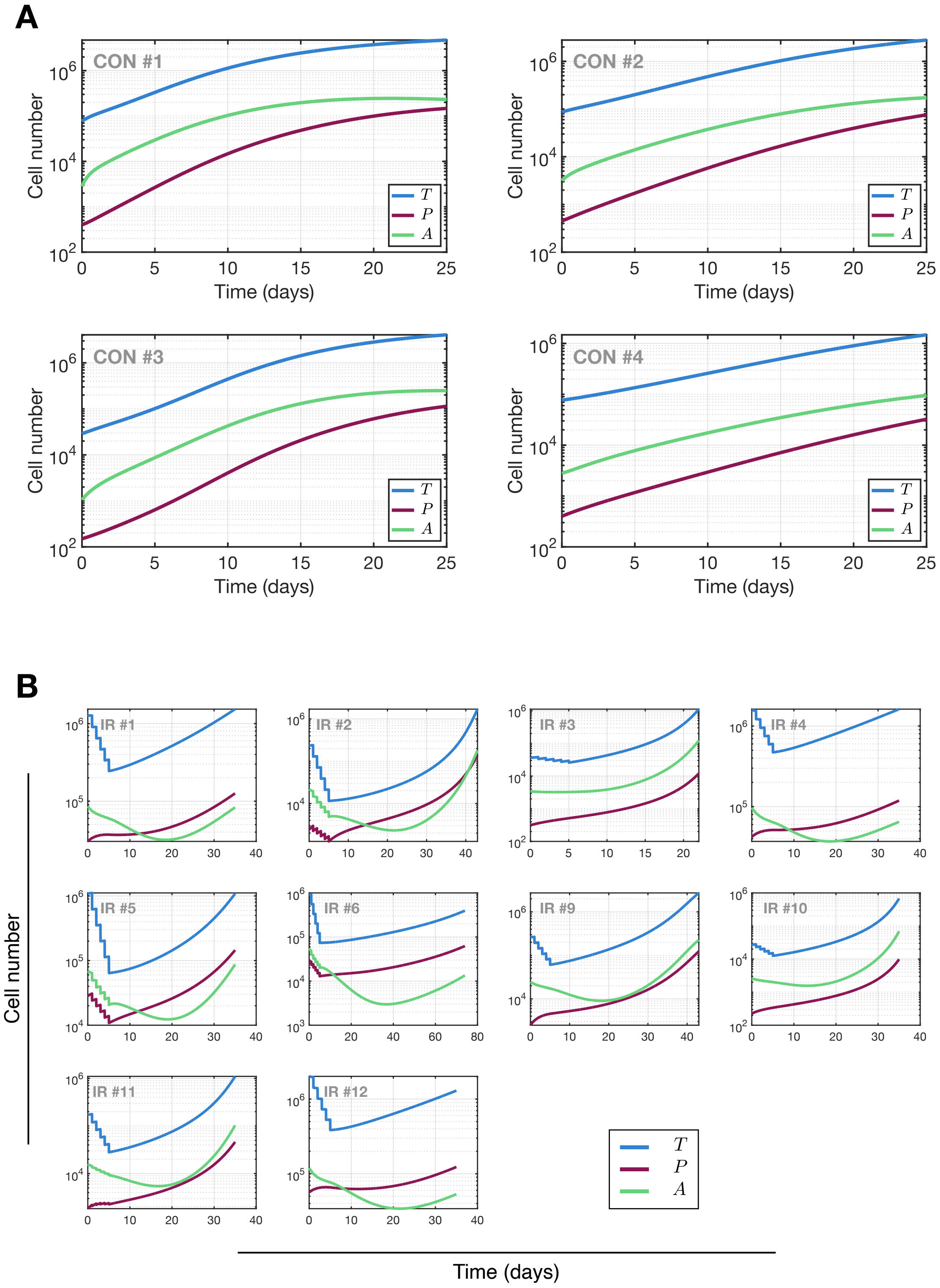


**Figure S4. Evolution of model variables.** Time evolution of the main populations in the model: glioma cells ($T$), anti-tumor macrophages ($A$), and pro-tumor macrophages ($P$). (A) shows the evolution in the control cases, while (B) shows the mice treated with 5 fractions of IR between day 1 and 6.

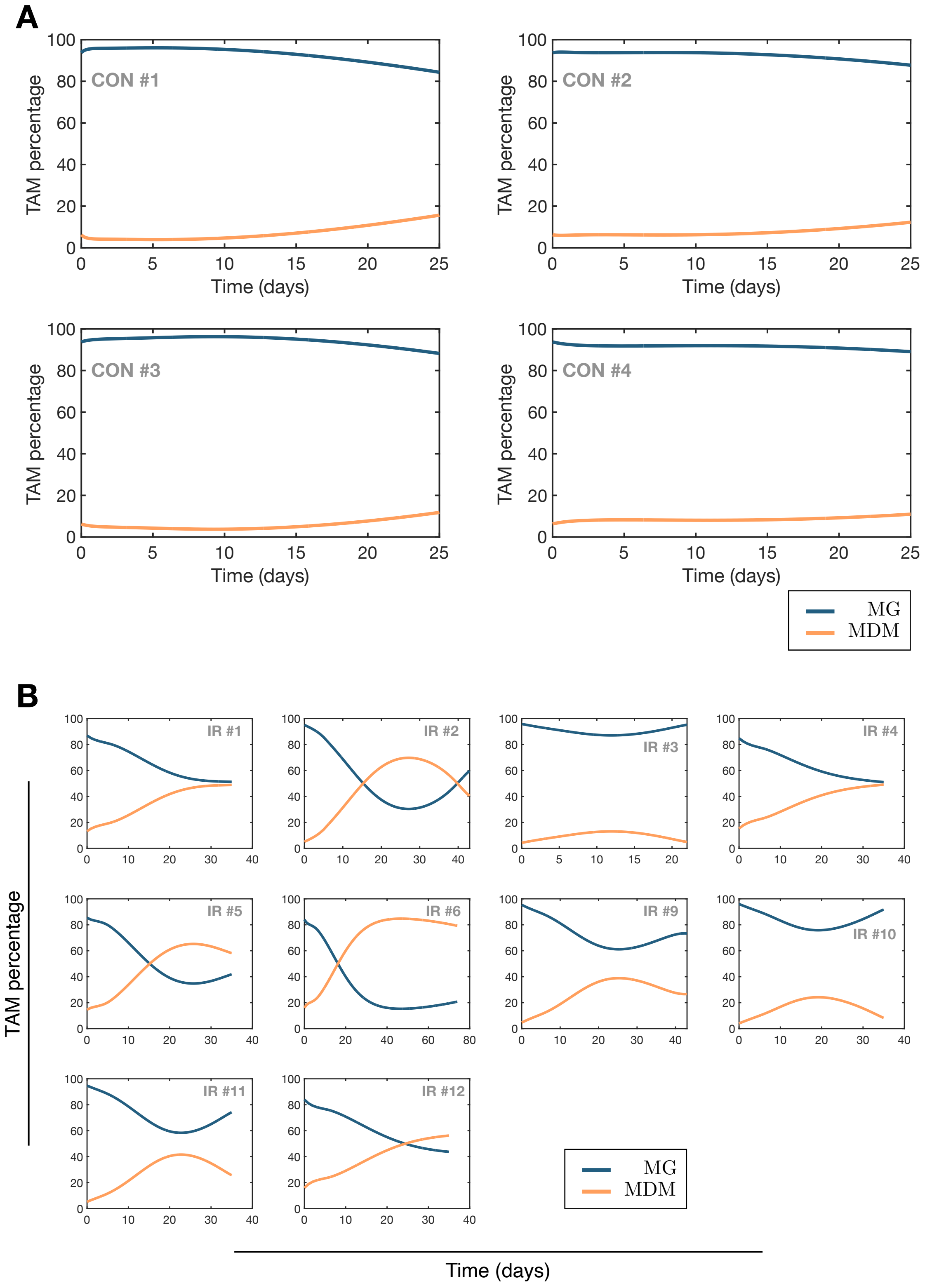


**Figure S5. Distribution of macrophages between MG and MDM.** The distribution of macrophages between resident microglia (MG) and infiltrating myeloid-derived macrophages (MDM). (A) control cases. (B) mice undergoing with 5 daily fractions of IR starting on day 1.

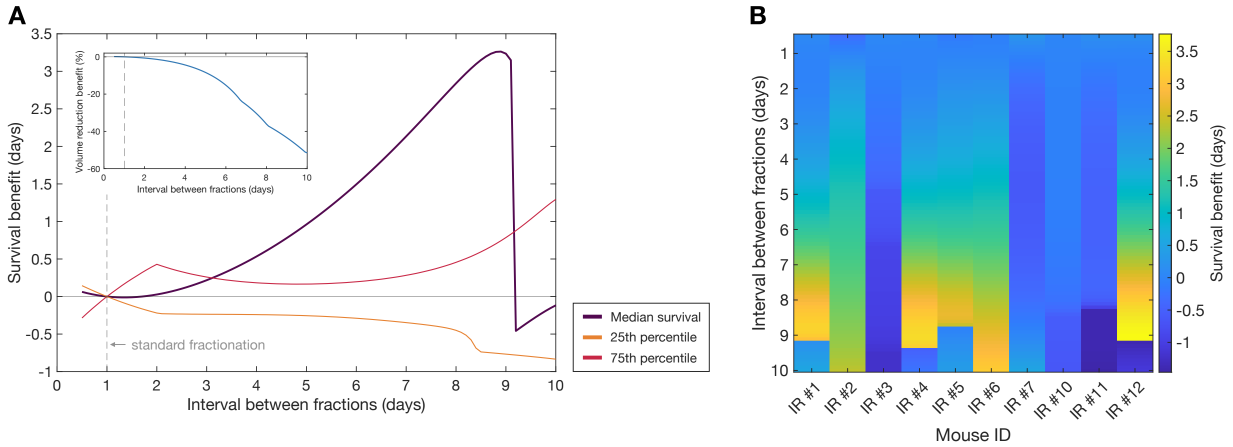


**Figure S6. Survival benefit of increasing the interval between IR fractions.** (A) Survival benefit in days of spacing out IR fractionations beyond the standard daily administration. The purple line represents the median gain, the maroon line shows the benefit at the 75th survival percentile (good survival subjects), and the orange line indicates the benefit (negative in this case, showing lower survival gains) for the 25th survival percentile (poor survival). For each studied fractionation interval, the inset shows the percentage of volume reduction lost due to the increased fractionation time. (B) Survival benefit (color) obtained by each individual mouse (x-axis) at each interval (y-axis).

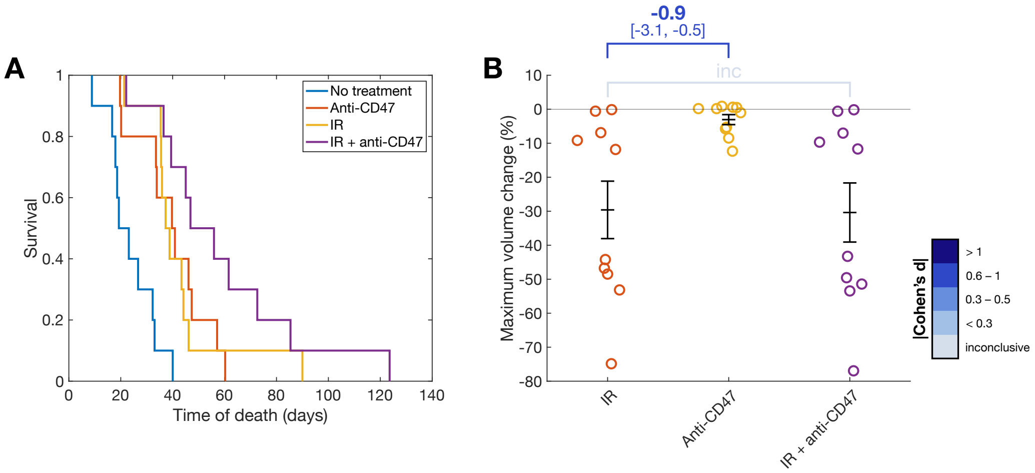


**Figure S7. Survival benefit of treatment with anti-CD47.** (A) Kaplan-Meier curves showing survival of mice digital twin under different treatments: no treatment (blue), anti-CD47 treatment (orange), IR (yellow), and IR + anti-CD47 (purple). (B) Maximum tumor volume change for each treatment. While anti-CD47 treatment alone provides some survival benefit, it is not effective in tumor debulking; graph shows means ± SEM and effect size is reported using Cohen's d, along with its confidence interval shown in square brackets..

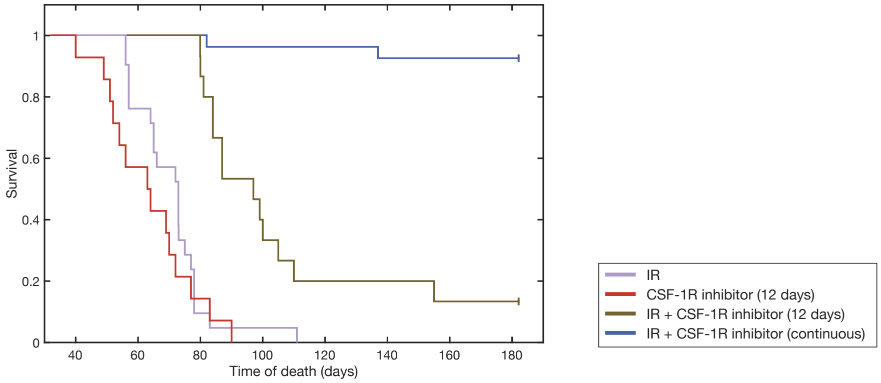


**Figure S8. Survival outcomes in PDG-Ink4a/ArfKO mice under CSF-1R inhibition.** Survival results from experiments on PDG-Ink4a/ArfKO mice treated with the following protocols: 5 daily fractions of IR alone, 12 days of CSF-1R inhibitor alone, combination of IR and 12 days of CSF-1R inhibitor, and 5 daily fractions of IR with continuous CSF-1R inhibitor treatment. Data sourced from ref. [2].

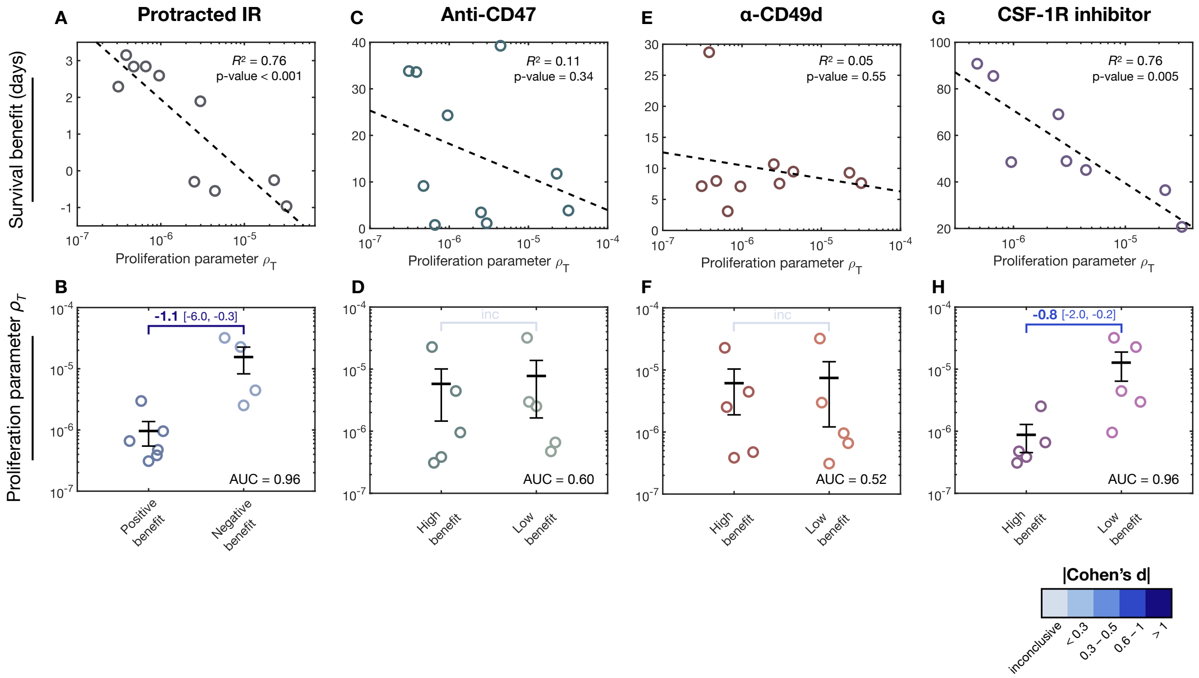


**Figure S9. Differential effect of the therapies for different proliferation rates.** The top row shows the linear correlation between the survival benefit of the different simulated treatments (structured in columns). The bottom row shows the values of the fitted tumor proliferation parameter $\rho_T$ for groups receiving a high and low benefit from the treatment.

### References


[1] F. Klemm, R. R. Maas, R. L. Bowman, M. Kornete, K. Soukup, S. Nassiri, J.-P. Brouland, C. A. Iacobuzio-Donahue, C. Brennan, V. Tabar, Interrogation of the microenvironmental landscape in brain tumors reveals disease-specific alterations of immune cells, Cell 181 (7) (2020) 1643–1660. doi:10.1016/j.cell.2020.05.007.

[2] L. Akkari, R. L. Bowman, J. Tessier, F. Klemm, S. M. Handgraaf, M. de Groot, D. F. Quail, L. Tillard, J. Gadiot, J. T. Huse, et al., Dynamic changes in glioma macrophage populations after radiotherapy reveal csf-1r inhibition as a strategy to overcome resistance, Science translational medicine 12 (552) (2020) eaaw7843.

[3] D. F. Quail, J. A. Joyce, The microenvironmental landscape of brain tumors, Cancer cell 31 (3) (2017) 326–341. doi:10.1016/j.ccell.2017.02.009.

[4] M. B. Graeber, B. W. Scheithauer, G. W. Kreutzberg, Microglia in brain tumors, Glia 40 (2) (2002) 252–259. doi:10.1002/glia.10147.

[5] D. H. Gutmann, H. Kettenmann, Microglia/brain macrophages as central drivers of brain tumor pathobiology, Neuron 104 (3) (2019) 442–449.

[6] S. M. Pyonteck, L. Akkari, A. J. Schuhmacher, R. L. Bowman, L. Sevenich, D. F. Quail, O. C. Olson, M. L. Quick, J. T. Huse, V. Teijeiro, et al., CSF-1R inhibition alters macrophage polarization and blocks glioma progression, Nature medicine 19 (10) (2013) 1264–1272. doi:10.1038/nm.3337.

[7] P. Schol, M. J. van Elsas, J. Middelburg, M. K. N. Twilhaar, T. van Hall, T. C. van der Sluis, S. H. van der Burg, Myeloid effector cells in cancer, Cancer Cell 42 (12) (2024) 1997–2014.

[8] W. Zhou, S. Q. Ke, Z. Huang, W. Flavahan, X. Fang, J. Paul, L. Wu, A. E. Sloan, R. E. McLendon, X. Li, et al., Periostin secreted by glioblastoma stem cells recruits m2 tumour-associated macrophages and promotes malignant growth, Nature cell biology 17 (2) (2015) 170–182. doi:10.1038/ncb3090.

[9] D. Hambardzumyan, D. H. Gutmann, H. Kettenmann, The role of microglia and macrophages in glioma maintenance and progression, Nature neuroscience 19 (1) (2016) 20–27.

[10] G. Fries, A. Perneczky, O. Kempski, Glioblastoma-associated circulating monocytes and the release of epidermal growth factor, Journal of neurosurgery 85 (4) (1996) 642–647.

[11] C. Huettner, S. Czub, S. Kerkau, W. Roggendorf, J.-C. Tonn, Interleukin 10 is expressed in human gliomas in vivo and increases glioma cell proliferation and motility in vitro., Anticancer research 17 (5A) (1997) 3217–3224.

[12] F. Khan, L. Pang, M. Dunterman, M. S. Lesniak, A. B. Heimberger, P. Chen, et al., Macrophages and microglia in glioblastoma: heterogeneity, plasticity, and therapy, The Journal of clinical investigation 133 (1) (2023).

[13] V. A. Kuznetsov, I. A. Makalkin, M. A. Taylor, A. S. Perelson, Nonlinear dynamics of immunogenic tumors: parameter estimation and global bifurcation analysis, Bulletin of mathematical biology 56 (2) (1994) 295–321.

[14] R. Eftimie, C. Barelle, Mathematical investigation of innate immune responses to lung cancer: The role of macrophages with mixed phenotypes, Journal of Theoretical Biology 524 (2021) 110739.

[15] S. Sabir, O. León-Triana, S. Serrano, R. Barrio, V. M. Pérez-García, Mathematical model of car t-cell therapy for a b-cell lymphoma lymph node, Bulletin of Mathematical Biology 87 (3) (2025) 1–33.

[16] J. H. Sampson, M. D. Gunn, P. E. Fecci, D. M. Ashley, Brain immunology and immunotherapy in brain tumours, Nature Reviews Cancer 20 (1) (2020) 12–25.

[17] B. Frey, S. Hehlgans, F. Rödel, U. S. Gaipl, Modulation of inflammation by low and high doses of ionizing radiation: Implications for benign and malign diseases, Cancer letters 368 (2) (2015) 230–237.

[18] S. Goswami, S. Anandhan, D. Raychaudhuri, P. Sharma, Myeloid cell-targeted therapies for solid tumours, Nature Reviews Immunology 23 (2) (2023) 106–120.

[19] S. Marelli, B. Sudret, UQLab: A Framework for Uncertainty Quantification in Matlab, American Society of Civil Engineers, 2011, pp. 2554–2563. arXiv:https://ascelibrary.org/doi/pdf/10.1061/9780784413609.257, doi:10.1061/9780784413609.257. URL https://ascelibrary.org/doi/abs/10.1061/9780784413609.257
